# A Missense Mutation in Human CHD4 Causes Ventricular Noncompaction by Repressing ADAMTS1-mediated Trabeculation Cessation

**DOI:** 10.1101/2022.09.12.507607

**Authors:** Wei Shi, Angel P. Scialdone, James I. Emerson, Liu Mei, Lauren K. Wasson, Haley A. Davies, Christine E. Seidman, Jonathan G. Seidman, Jeanette G. Cook, Frank L. Conlon

## Abstract

**Background:** Left ventricular noncompaction (LVNC) is a prevalent cardiomyopathy associated with excessive trabeculation and thin compact myocardium. Patients with LVNC are vulnerable to cardiac dysfunction and at high risk of sudden death. Although sporadic and inherited mutations in cardiac genes are implicated in LVNC, understanding of the mechanisms responsible for human LVNC is limited.

**Methods:** We screened the complete exome sequence database of the Pediatrics Cardiac Genomics Consortium and identified a cohort with a *de novo* chromodomain helicase DNA-binding protein 4 (CHD4) proband, CHD4^M202I^, with congenital heart defects. We engineered a patient-specific model of CHD4^M202I^ (mouse CHD4^M195I^). Histological analysis, immunohistochemistry, flow cytometry, transmission electron microscopy, and echocardiography were used to analyze cardiac anatomy and function. *Ex vivo* culture, immunopurification coupled with mass spectrometry, transcriptional profiling, and chromatin immunoprecipitation were performed to deduce the mechanism of CHD4^M195I^-mediated ventricular wall defects.

**Results:** *CHD4^M195I/M195I^* mice developed biventricular hypertrabeculaion and noncompaction and died at birth. Proliferation of cardiomyocytes was significantly increased in *CHD4^M195I^* hearts, and the excessive trabeculation was associated with accumulation of extracellular matrix (ECM) proteins and a reduction of ADAMTS1, an ECM protease. We rescued the hyperproliferation and hypertrabeculation defects in *CHD4^M195I^* hearts by administration of ADAMTS1. Mechanistically, the CHD4^M195I^ protein showed augmented affinity to endocardial BRG1. This enhanced affinity resulted in failure of derepression of *Adamts1* transcription such that ADAMTS1-mediated trabeculation termination was impaired.

**Conclusions:** Our study reveals how a single mutation in the chromatin remodeler CHD4, in mice or humans, modulates ventricular chamber maturation and that cardiac defects associated with the missense mutation CHD4^M195I^ can be attenuated by the administration of ADAMTS1.

**Clinical Perspective:** *What Is New?:*

- A patient-specific mouse model of *CHD4^M202I^* develops ventricular hypertrabeculation and dies at birth.
- Proliferation of cardiomyocytes is significantly enhanced in *CHD4^M195^*^I^ mice.
- ADAMTS1 is significantly downregulated in *CHD4^M195I^* mice.
- Close interaction between CHD4^M195I^ and BRG1 robustly and continuously represses *Adamts1* transcription, which impairs ADAMTS1-mediated termination of trabeculation in the developing mutant heart.

*What Are the Clinical Implications?:*

- This study provides a unique mouse model of ventricular noncompaction cardiomyopathy that faithfully reflects human patients’ genetic condition without disturbing the target gene’s expression and localization.
- Transcriptional repression of ECM protease ADAMTS1 by CHD4-BRG1 interaction is detrimental to ventricular wall maturation; maintaining appropriate ADAMTS1 levels in the heart could be a promising therapeutic approach for treating ventricular noncompaction cardiomyopathy.

## Introduction

Left ventricular noncompaction (LVNC), the third most prevalent cardiomyopathy, is characterized by excessive ventricular trabeculae (noncompaction), a thin myocardium, and deep intertrabecular recesses among the trabeculae. Left ventricular noncompaction mainly affects the left ventricle, but isolated right ventricular or biventricular noncompaction has also been reported^1–5^. Left ventricular noncompaction can be asymptomatic, but patients with LVNC are at high risk to develop heart failure and sudden death^6^. Genetic screening of individual LVNC patients has identified mutations in genes encoding transcription factors, sarcomere, nuclear membrane, RNA-binding proteins, and ion channel proteins^7–10^; however, a significant portion of LVNC cases are of unknown etiology^11^. Moreover, identification of the molecular and cellular mechanisms that orchestrate the pathogenesis of LVNC is compromised by the absence of a genetic model that faithfully recapitulates the conditions of patients.

As a structural cardiac defect, LVNC involves ventricular chamber maturation, during which ventricular trabeculation and compaction are the most important morphogenetic events^12^. Trabeculation is initiated by myocardial sprouting that extends from the compact monolayer and protrudes into the extracellular matrix (ECM), also referred to as the cardiac jelly at approximately E8.5 in mouse and approximately day 28 in human^13^. Trabecular myocardium grows by proliferation of cardiomyocytes at early developmental stages. By E13.5 in mouse and approximately day 40 in human, the trabeculae undergo significant compaction and form a thickened, tightly packed ventricular wall with a relatively smooth inner surface^12^. Persistent trabeculation and a reduced level of compaction^6, 14, 15^ typically characterize LVNC.

Abnormal development of the ventricular chambers of the heart is the basis of a significant number of congenital heart defects. Therefore, a mechanistic understanding of cardiac compaction is crucial for improving treatment of structural heart disease. Large, multi-component complexes that modify chromatin control cardiac gene expression. Prominent among these complexes is the Nucleosome Remodeling and Deacetylase (NuRD) complex^16–18^. The NuRD complex is essential for many developmental events, including ensuring proper timing of the switch from stem cell lineages to differentiated cell types, maintaining cell differentiation, and activating DNA damage response pathways^19–24^. Mutations in chromodomain helicase DNA-binding protein 4 (CHD4), the core catalytic component of NuRD, lead to congenital cardiac malformation, including atrial and ventricular septal defects^25^. The CHD4 component is essential for cardiac development, and it represses inappropriate expression of the skeletal and smooth muscle programs in the developing heart^26–28^. Nonetheless, neither CHD4 nor the NuRD complex can directly bind DNA. Instead, we have shown that CHD4 is recruited to target cardiac genes by interaction with the essential cardiac transcription factors GATA4, NKX2-5, and TBX5^29^.

Although CHD4 is crucial for heart development, and its disease relevance is well established, many questions remain regarding the mechanism of CHD4’s function. Critically, we do not understand how a single missense mutation in CHD4, a broadly expressed catalytic unit of the NuRD complex, leads to cardiac-specific disease in human. To determine the basis of CHD4 mutations in human congenital heart disease, we screened the complete exome sequence database of the Pediatrics Cardiac Genomics Consortium^30, 31^ and identified a cohort with a *de novo* CHD4 proband, CHD4^M202I^, with congenital heart defects. Using CRISPR/CAS9, we generated a mouse patient-specific model of CHD4^M202I^ (mouse CHD4^M195I^). We demonstrate that *CHD4^M195I/M195I^* mice displayed critical aspects of human LVNC. *CHD4^M195I/M195I^* hearts failed to undergo compaction, and cardiomyocytes were retained in an immature proliferating state through birth. The LVNC phenotype was associated with an increase in extracellular matrix proteins and a marked reduction in the ECM protease ADAMTS1. We demonstrate that *Adamts1* is a direct target of CHD4 and that the exogenous administration of ADAMTS1 or the introduction of a pharmacological compound that increases ADAMTS1 expression rescues key facets of LVNC *ex vivo* and in utero in *CHD4^M195I/M195I^* mice. Together, we have revealed a mechanism of how CHD4 may cause congenital heart diseases, and we present prospects for treatment.

## Methods

### Ethics Statement

The Institutional Animal Care and Use Committee at the University of North Carolina at Chapel Hill approved all animal experiments, which conformed to the Guide for the Care and Use of Laboratory Animals. An expanded Methods section is available in the Supplemental Material. On reasonable request, the data, analytic methods, and study materials will be made available to other researchers for purposes of reproducing the results or replicating the procedures.

## Results

### Generation of a patient-specific mouse model of *CHD4^M202I^*

To define the mechanisms by which mutations in CHD4 lead to cardiac disease, we screened the complete exome sequence database of the Pediatrics Cardiac Genomics Consortium^31^ and identified a cohort with a *de novo* CHD4 proband CHD4^M202I^. This proband was associated with ventricular septal defects and conotruncal abnormalities (Figure 1A). Surprisingly, CHD4^M202I^ mapped to a highly evolutionarily conserved region (Figure S1A) of CHD4 of unknown biological function^32–34^. To determine the mechanism by which CHD4^M202I^ causes cardiac disease, we used a CRISPR/CAS9 gene-editing system to engineer a mouse patient-specific model of CHD4^M202I^ (mouse CHD4^M195I^; Figure S1B). Founding males were bred to wild-type C57BL/6J females, and the genotypes of the founding males and all F2 offspring were confirmed by PCR genotyping and sequencing (Figure S1C and S1D). Immunoblot analyses and immunohistochemistry at E18.5 showed that intact CHD4^M195I^ protein was present in homozygous *CHD4^M195I/M195I^* hearts, at endogenous levels and localized to the nucleus (Figure S1E and S1F).

**Figure 1.**
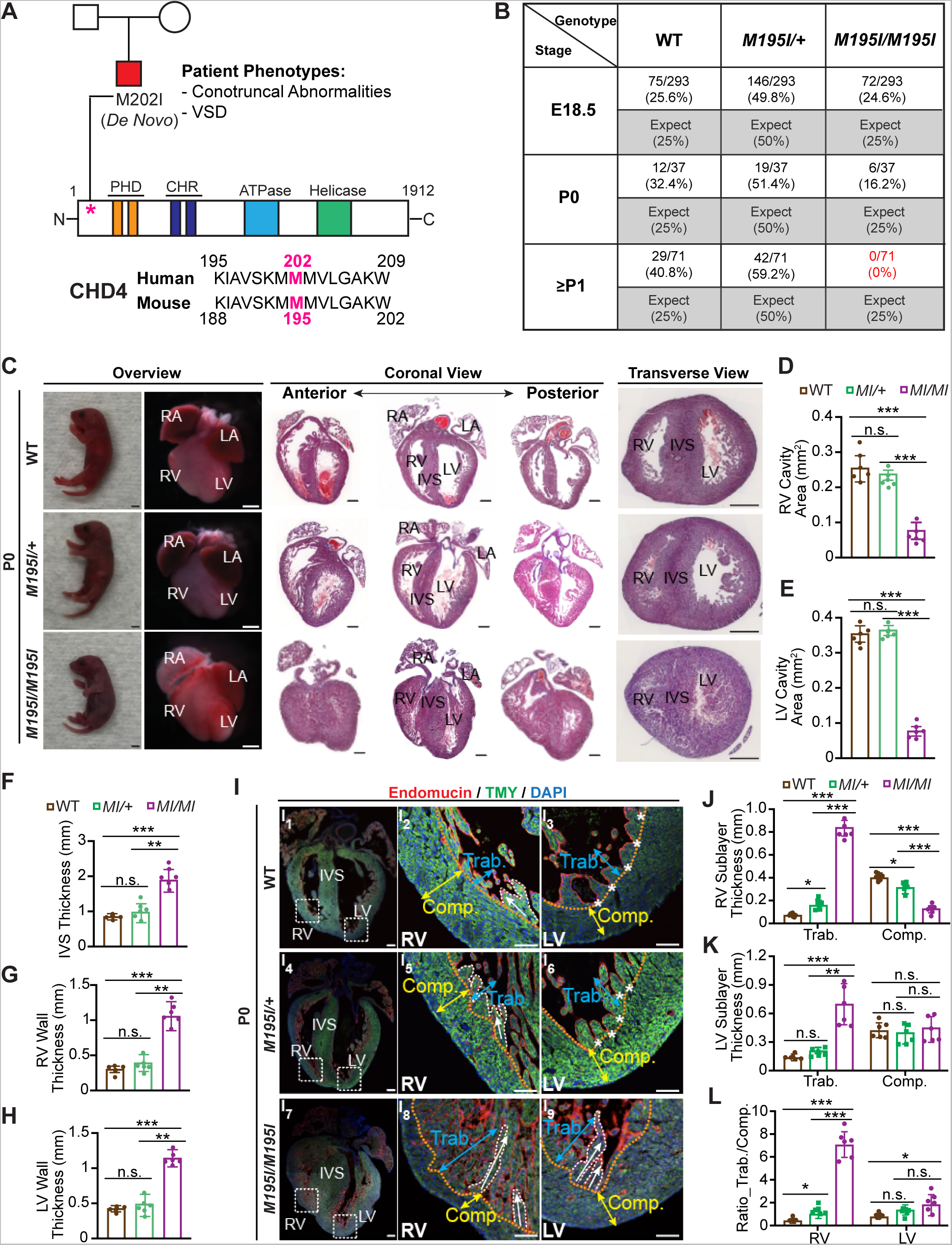
Homozygous *CHD4^M195I^* mutant neonatal mice develop ventricular noncompaction and die at birth. **A**, Top, CHD4^M202I^ pedigree (squares, male; circle, female; VSD, ventricular septal defect) and schematic of CHD4 protein domains; PHD, plant homeodomain; CHR, chromodomain. Bottom, protein sequence alignment of human and mouse CHD4 orthologs. **B**, Table of expected and observed Mendelian ratios of offspring at indicated for developmental stages from the intercross of *CHD4^M195I/+^* mice. E, embryonic day; P, postnatal day. **C**, Representative P0 neonates, whole mount and H&E-stained paraffin coronal and transverse sections of P0 *WT*, heterozygote (*CHD4^M195I/+^*), and homozygote (*CHD4^M195I/M195I^*) mouse hearts. Scale bars, 2 mm (overview of neonates) and 0.5 mm (overview of whole mount and H&E stains). RA: right atrium; RV: right ventricle; LA: left atrium; LV: left ventricle; IVS: interventricular septum. **D**-**H**, Quantification of right (**D**) and left (**E**) ventricular cavity area, IVS (**F**), RV wall (**G**), and LV wall (**H**) thickness on P0 WT, *CHD4^M195I/M195I^*, and *CHD4^M195I/M195I^* mouse heart sections (n=6 independent hearts per genotype). **I**, Representative images of immunofluorescent (Endomucin, Tropomyosin (TMY), and DAPI-stained sections from P0 mouse hearts. The approximate boundaries between compact myocardium (Comp.; indicated by yellow double-headed arrows) and trabecular myocardium (Trab.; indicated by blue double-headed arrows) are indicated by orange dashed lines. Trabeculae are delineated with white dashed curve, and fused trabeculae are indicated by white asterisks. White arrows indicate the direction of trabecular projections. Scale bars, 0.5 mm (overview of whole heart) and 0.2 mm (magnified views on RV and LV). **J** and **K**, Measurements of RV (**J**) and LV (**K**) sublayer thickness on P0 mouse hearts (n=6 individual hearts per genotype). **L**, Ratio of trabecular layer thickness to compact layer thickness in (**J**) and (**K**). Data in **D**-**H** and **J-L**, are represented as mean ± SEM. Statistical significance was determined with one-way ANOVA (n.s., not significant (P>0.05); \**P*<0.05, \*\**P*<0.01, \*\*\**P*<0.001). *M195I/M195I* or *MI/MI*: *CHD4^M195I/M195I^*, same with all main and supplemental figures.

### Neonatal homozygous *CHD4^M195I/M195I^* develop noncompaction cardiomyopathy

Heterozygous *CHD4^M195I/+^* mice were viable, fertile, recovered at expected Mendelian ratios, and exhibited no overt phenotypic abnormalities (Figure 1B and Figure S2). Conversely, homozygous *CHD4^M195I/M195I^* mice died at or shortly after birth, with none of the *CHD4^M195I/M195I^* mice surviving more than two days (Figure 1B). Histological examination at postnatal (P) 0 revealed that *CHD4^M195I/M195I^* mouse hearts had a dramatically reduced ventricular cavity with a concomitant increase in the thickness of the ventricular walls and septa compared with WT or *CHD4^M195I/+^* (Figure 1C-1H).

Left ventricular noncompaction (LVNC) is a cardiac disease associated with an increase in cardiac failure in children and adults^35, 36^. In addition, LVNC is associated with hypertrabeculation of the myocardium of the left ventricle (LV), and occasionally the right ventricle (RV)^1–5^. Therefore, to assess the compact and trabecular layers of *CHD4^M195I/M195I^* embryos, we conducted immunofluorescence with Endomucin (red) and Tropomyosin (green) antibodies to delineate chamber endocardium and myocardium, respectively (Figure 1I and Figure S3). In WT and *CHD4^M195I/+^* hearts, the trabeculae were folded and oriented parallel to the free wall of the right ventricle (white arrows, Figure 1I_2_ and I_5_). Moreover, the trabeculae assimilated into the compact zones clearing the ventricular lumen (asterisks, Figure 1I_3_-1I_6_). In contrast, and consistent with LVNC phenotype^6, 14, 15^, the trabeculae in the *CHD4^M195I/M195I^* hearts remained straight and protruded perpendicularly to the compact zones and extended across the lumen (white arrows, Figure 1I_8_ and 1I_9_). By examining the ratio of thickness of trabecular-to-compact layer, we found severe RV and modest LV noncompaction in *CHD4^M195I/M195I^* hearts. In contrast, *CHD4^M195I/+^* hearts displayed RV but not LV noncompaction (Figure 1J-1L). Together, these findings demonstrated that *CHD4^M195I^* is associated with critical aspects of the LVNC phenotype.

### Ventricular noncompaction in *CHD4^M195I/M195I^* hearts

To define the onset of noncompaction in *CHD4^M195I/M195I^* embryos, we conducted histological analysis on *CHD4^M195I/M195^* and WT embryonic hearts from E9.5 through E18.5. At E9.5, LV and RV in WT embryos displayed trabeculae as single sprouts with few interconnections (Figure 2A_1_-2A_3_). In contrast, trabeculae in the *CHD4^M195I/M195I^* hearts were longer, they protruded into the ventricular cavities, and they formed a ventricular mesh network (Figure 2A_4_-2A_6_ and 2B). At this stage, alternations in morphology of *CHD4^M195I/M195I^* hearts appeared to be confined to the trabecular layer, and we did not observe a difference in thickness of the compact layer in either the RV or LV (Figure 2C). However, by E12.5, in *CHD4^M195I/M195I^* hearts we found a thinning of the compact layer in addition to excess trabeculae (Figure S4A-S4C).

**Figure 2.**
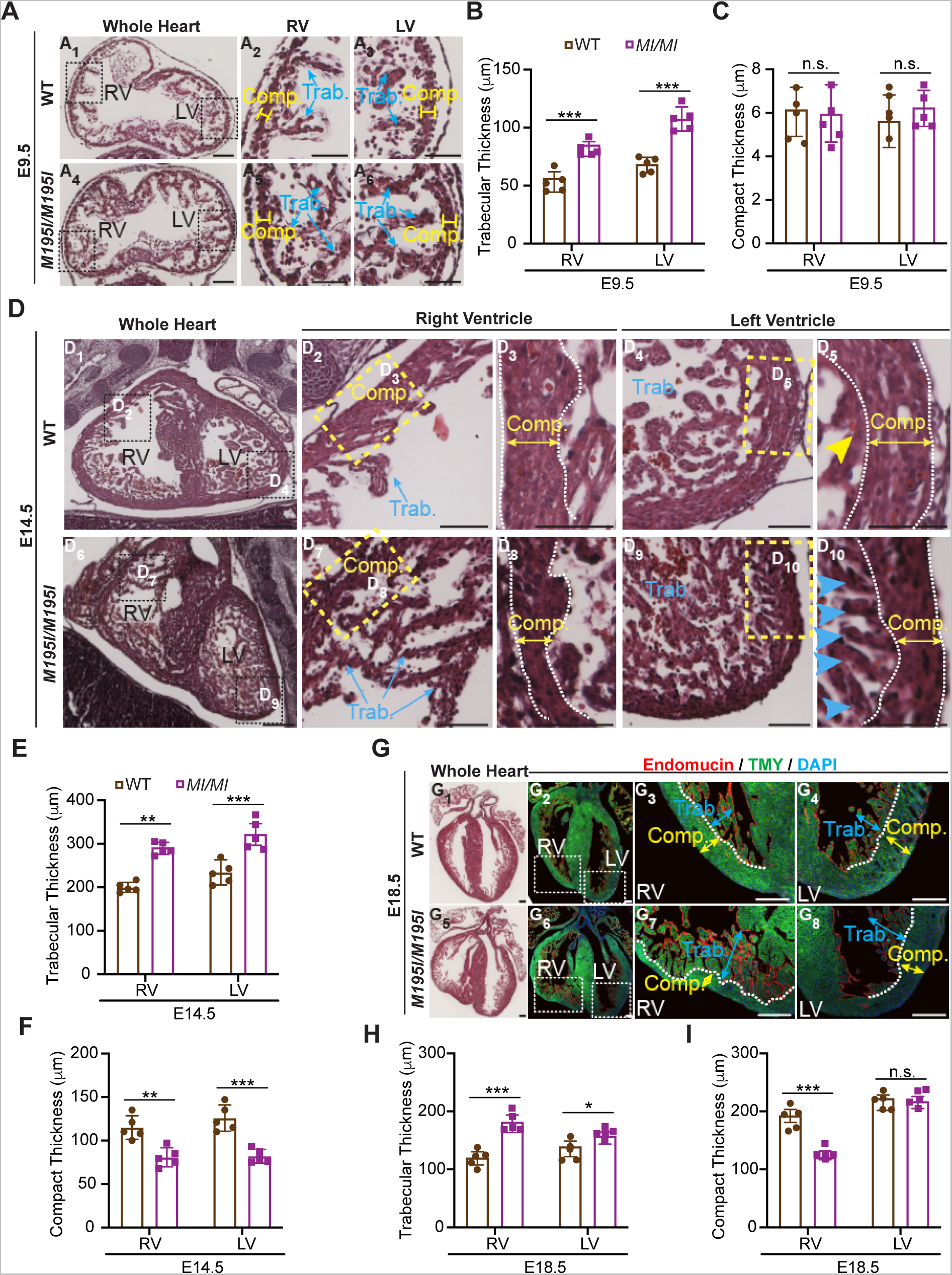
Ventricular noncompaction in homozygous *CHD4^M195I/M195I^* hearts dynamically proceeds over developmental stages. **A** and **D**, Representative images of H&E-stained paraffin sections of WT and *CHD4^M195I/M195I^* hearts at E9.5 (**A**) and E14.5. (**D**), trabeculae assimilated into the compact myocardium are indicated by yellow arrowhead (**D_5_**) and non-assimilated trabeculae are indicated by blue arrowheads (**D_10_**). **G**, Representative images of H&E-stained (**G_1_** and **G_5_**) and immunofluorescent-(Endomucin, TMY, and DAPI; **G_2_-G_4_** and **G_6_-G_8_**) stained paraffin sections of WT and *CHD4^M195I/M195I^* hearts at E18.5; the approximal boundaries between compact myocardium (Comp.; indicated by yellow double-headed arrows) and trabecular myocardium (Trab.; indicated by blue double-headed arrows) are indicated by orange dashed lines. Scale bars, 50 μm (**A**), 250 μm (**D**), and 200 μm(**G**). **B**, **C**, **E**, **F**, **H**, and **I**, Measurements of sublayer thicknesses of WT and *CHD4^M195I/M195I^*heart sections from E9.5 (**B** and **C**), E14.5 (**E** and **F**), and E18.5 (**H** and **I**). Data in **B**, **C**, **E**, **F**, **H**, and **I** are represented as mean ± SEM, n=5 for each genotype per stage. Statistical significance was determined with two-tailed unpaired Student’s *t* test (n.s., not significant (P>0.05); \**P*<0.05, \*\**P*<0.01, \*\*\**P*<0.001).

By E14.5, in WT hearts, portions of the trabeculae were assimilated into an organized compact zone (Figure 2D_1_-2D_5_), whereas in *CHD4^M195I/M195I^* hearts the compact layer was thinner, the trabeculae were thicker, and we observed a failure of the trabeculae to fold (Figure 2D_6_-2D_10_, 2E and 2F). These phenotypes persisted in E16.5 (Figure S4D-S4F) and E18.5 (Figure 2G-2I). However, the thickness of the compact zones in LV was not significantly different compared with WT hearts at these stages (Figure 2I and Figure S4F) and at P0 (Figure 1K and 1L). These findings demonstrated that *CHD4^M195I/M195I^* hearts underwent dynamic remodeling leading to an LVNC phenotype and death at or shortly after birth.

### Impaired cardiac function in *CHD4^M195I/M195I^* mice

To delineate the physiological consequences of *CHD4^M195I^*, we performed ultrasound pulsed wave (PW) Doppler on E18.5 embryos in utero without surgical manipulation of the dam or embryos^26, 37, 38^. This approach enabled us to measure the effect of biventricular noncompaction on cardiac function in the developing heart. Short axis M-mode echocardiograms showed little ventricular wall movement in the *CHD4^M195I/M195I^* (Figure 3A), which confirmed severe cardiac contractile dysfunction and consequently reduced heart rate (Figure 3B), left ventricular ejection fraction (LV EF; Figure 3C), left ventricular fractional shortening (LV FS; Figure 3D), right ventricular end-systolic and diastolic volume (RV Vol;s, Figure 3E; RV Vol;d, Figure 3F). No significant changes were identified in the *CHD4^M195I/+^* embryos or adults (Figure 3A and Figure S5). These studies demonstrated that *CHD4^M195I/M195I^* physiologically models LVNC.

**Figure 3.**
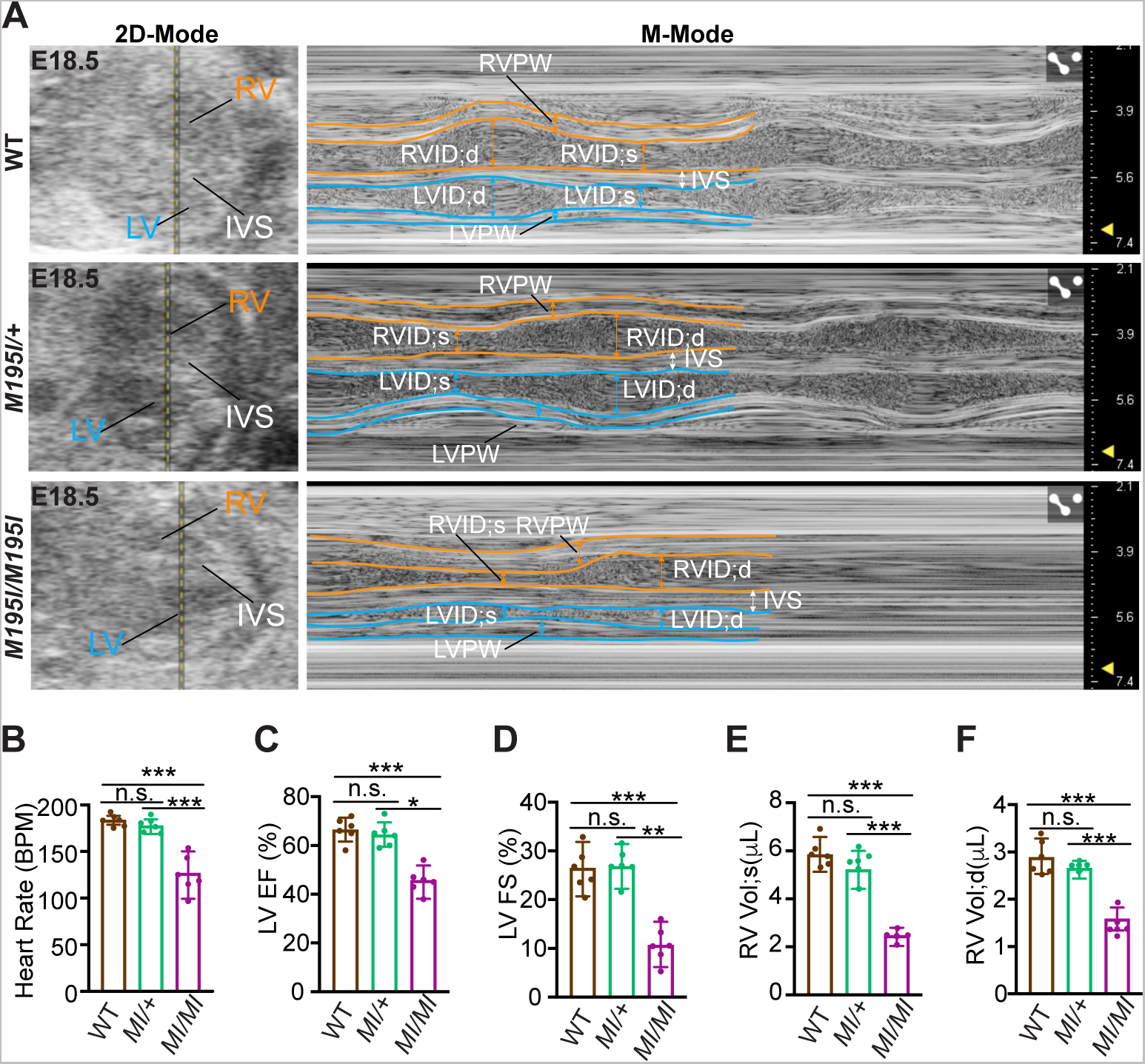
Impaired cardiac function in homozygous *CHD4^M195I/M195I^ mice*. **A**, In utero echocardiography analysis of representative E18.5 WT, *CHD4^M195I/+^,* and *CHD4^M195I/M195I^* embryonic hearts. Representative images showing 2D-Mode (left) and M-Mode (right) of the echocardiography measurement. RV, right ventricle; LV, left ventricle; IVS, interventricular septum; RVPW, right ventricular posterior wall; RVID;d, end-diastolic right ventricular internal diameter; RVID;s, end-systolic right ventricular internal diameter; LVID;d, end-diastolic left ventricular internal diameter; LVID;s, end-systolic left ventricular internal diameter; LVPW, left ventricular posterior wall. **B** through **F**, Quantification of heart rate (**B**), LV ejection fraction (LV EF, **C**), LV fractional shortening (LV FS, **D**), end-systolic right ventricular volume (RV Vol;s, **E**), and end-diastolic right ventricular volume (RV Vol;d, **F**). All data in **B** through **F** are represented as mean ± SEM, n=6 embryos per genotype from four pregnant female mice. Statistical significance was determined with one-way ANOVA (n.s., not significant (P>0.05); \**P*<0.05, \*\**P*<0.01, \*\*\**P*<0.001).

### Hypertrabeculation is accompanied by over-proliferation of cardiomyocytes

We next queried whether hypertrabeculation in *CHD4^M195I/M195I^* hearts was a result of the inappropriate proliferation of cardiomyocytes. Results revealed a significant increase in the co-expression of Ki67, a marker of proliferation and the cardiomyocyte marker TMY in *CHD4^M195I/M195I^* P0 mice (Figure 4A-4D). Consistently, quantification of dissociated cardiomyocytes from P0 hearts confirmed an increase in the number of cardiomyocytes in *CHD4^M195I/M195I^* hearts compared with WT (Figure S6A). Correspondingly, we found an increase in cardiomyocyte proliferation in vivo as measured by an increase in EdU incorporation (Figure 4E and 4F) and Ki67/TMY co-staining at E18.5 (Figure S6B and S6C).

**Figure 4.**
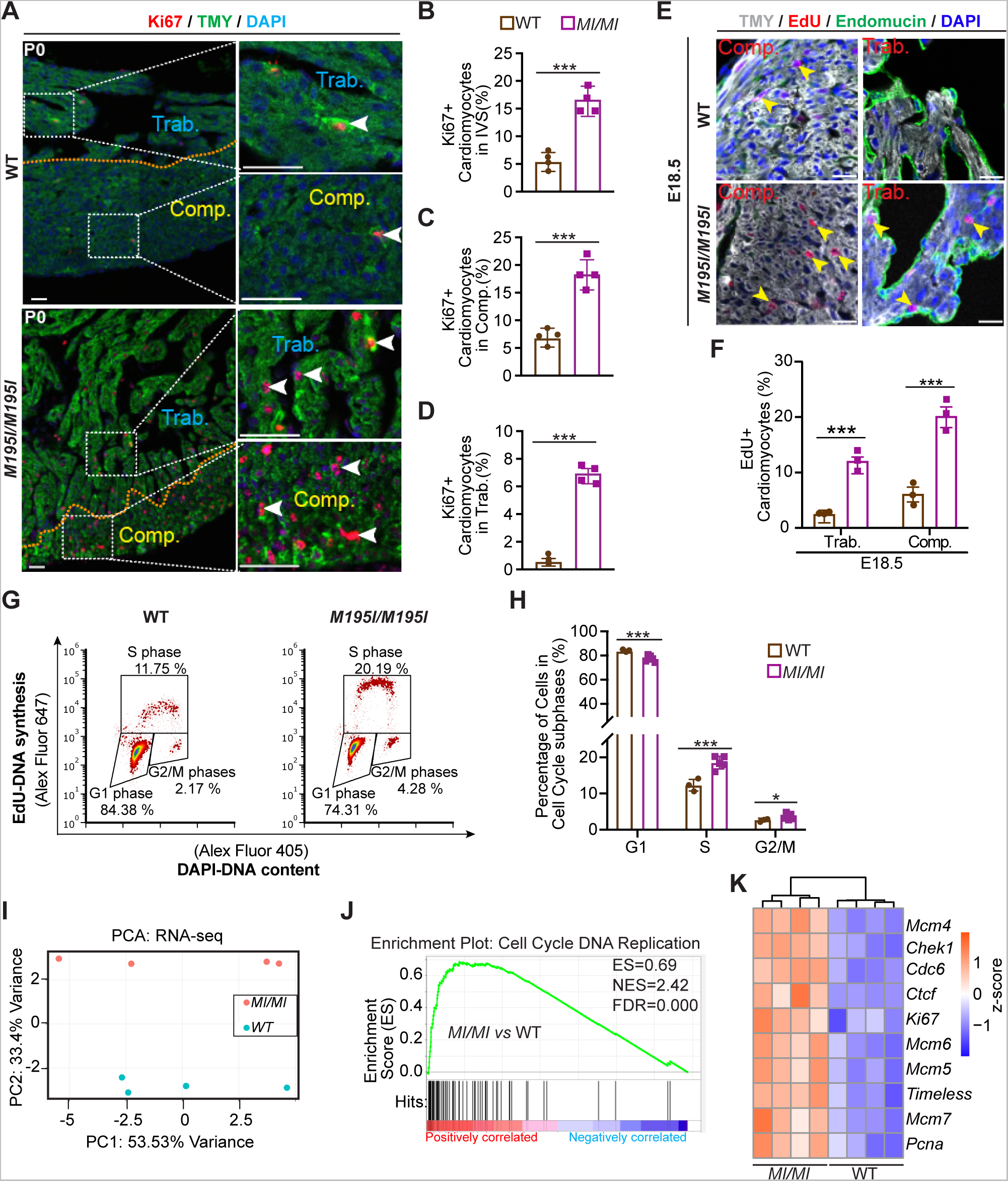
Cardiomyocyte proliferation is increased in *CHD4^M195I/M195I^*. **A**, Representative images of immunofluorescent (Ki67, TMY, and DAPI) stained paraffin sections from P0 WT and *CHD4^M195I/M195I^* mouse hearts. The approximate boundaries between compact myocardium (Comp.) and trabecular myocardium (Trab.) are indicated by orange dashed lines. Proliferating cardiomyocytes (Ki67^+^/TMY^+^) are indicated by white arrowheads. Scale bars: 50 μm. **B** through **D**, Quantification of proliferating cardiomyocytes ratios (Ki67^+^/total cardiomyocytes) in the interventricular septum (IVS, **B**), compact myocardium (Comp., **C**), and trabecular myocardium (Trab., **D**) (n=4 P0 hearts per genotype). **E**, Representative images of immunofluorescent (TMY, EdU, and DAPI) stained paraffin sections from E18.5 WT and *CHD4^M195I/M195I^* mouse hearts. Proliferating cardiomyocytes are indicated by yellow arrowheads. Scale bars, 20 μm. **F**, Quantification of proliferating cardiomyocytes ratio (EdU^+^/total cardiomyocytes) in E18.5 WT and *CHD4^M195I/M195I^* hearts (n=3 per genotype). **G**, Flow cytometry analysis for the proliferating cardiomyocytes derived from E18.5 *CHD4^M195I/M195I^* (n=5) and littermate WT (n=3) embryos. **H**, Quantification for the percent of cardiomyocytes at each cell cycle phase. **I**, Principal component analysis (PCA) plot of gene expression data from RNA-seq of four replicates for each genotype (WT and *CHD4^M195I/M195I^*). **J**, Gene set enrichment analysis (GSEA) showing the upregulated gene set of Cell Cycle DNA Replication in *CHD4^M195I/M195I^*. ES, enrichment score; NES, normalized enrichment score; FDR, false discovery rate. **K**, Heatmap of representative misregulated genes involved in **J**. Color bar: z-score. Data in **B** through **D**, **F** and **H** are represented as mean ± SEM, evaluated by two-tailed unpaired Student’s *t* test (n.s., not significant (P>0.05), \**P*<0.05, \*\**P*<0.01, \*\*\**P*<0.001).

To confirm and extend these findings, we conducted cell cycle profiling of cardiomyocytes derived from *CHD4^M195I/M195I^* and WT mice that had been labeled with EdU (Figure S7). We found a significant decrease in *CHD4^M195I/M195I^* (76.2%) compared with WT (84.3%) cardiomyocytes in G1 phase and a concomitant increase of cardiomyocytes in S phase (19.8% in *CHD4^M195I/M195I^* versus 12.3% in WT) and an increase in G2/M phases (3.8% in *CHD4^M195I/M195I^* versus 2.2% in WT) (Figure 4G and 4H). In sum, *CHD4^M195I/M195I^* cardiomyocytes displayed an increased and prolonged state of proliferation.

We further found that cardiomyocyte proliferation and hypertrabeculation were not associated with cardiomyocyte hypertrophy. Wheat Germ Agglutinin and Tropomyosin (TMY) double immunostaining (Figure S8A) on sections of P0 hearts did not show a significant difference in cardiomyocyte size between the WT and *CHD4^M195I/M195I^* hearts (Figure S8B). These results indicated that the hypertrabeculation was caused by over-proliferation of cardiomyocytes, and the overgrowth of the trabecular layer at the expense of the compact layer.

### CHD4^M195I^ leads to upregulation of cell cycle pathways

Because the phenotype and timing of *CHD4^M195I/M195I^* is significantly different from cardiac *Chd4* null mutations^26, 28^, we hypothesized that CHD4^M195I^ regulates a distinct transcriptional network essential for cardiac development and function. To test this hypothesis, we performed RNA-sequencing (RNA-seq) analyses on E18.5 WT and *CHD4^M195I/M195I^* hearts and identified 323 genes that were differentially expressed [adjusted P value <0.05, |log2(Fold Change)| > 0.5], 113 upregulated and 210 downregulated in *CHD4^M195I/M195 I^*(Figure 4I and Figure S9A). Consistent with our in vivo analyses at P0, Gene Set Enrichment Analysis (GSEA) revealed the overrepresented classifications in the *CHD4^M195I/M195I^* hearts to be those of cell cycle-related biological processes, including DNA replication initiation, cell cycle checkpoint control, nucleosome assembly, and cytokinesis (Figure 4J and Figure S9B). Representative differential genes included the core subunits of the DNA helicases *Mcm*(*4*-*7*), which are essential for DNA replication^39^ (Figure 4K). Thus, *CHD4^M195I^* was associated with an increase in components of the cell cycle and a related increase in cardiomyocyte proliferation.

### Cardiomyocytes in *CHD4^M195I/M195I^* are in an immature state

In mice at E18.5, cardiomyocytes initiate multiple adaptations for heart to transit from fetal to adult states^40–42^. In line with the higher cardiomyocyte proliferation activity of *CHD4^M195I/M195I^* hearts (Figure 4), our GSEA classification also revealed that the downregulated genes in *CHD4^M195I/M195I^* hearts were significantly enriched in differentiation functions, such as muscle cell development, sarcomere organization, and myofibril assembly (Figure 5A and Figure S6C). Myosin heavy chain 6 (*Myh6*), a dominant adult sarcomeric isoform in rodents^43, 44^, was among the representative downregulated genes in *CHD4^M195I/M195I^* hearts (Figure 5B).

**Figure 5.**
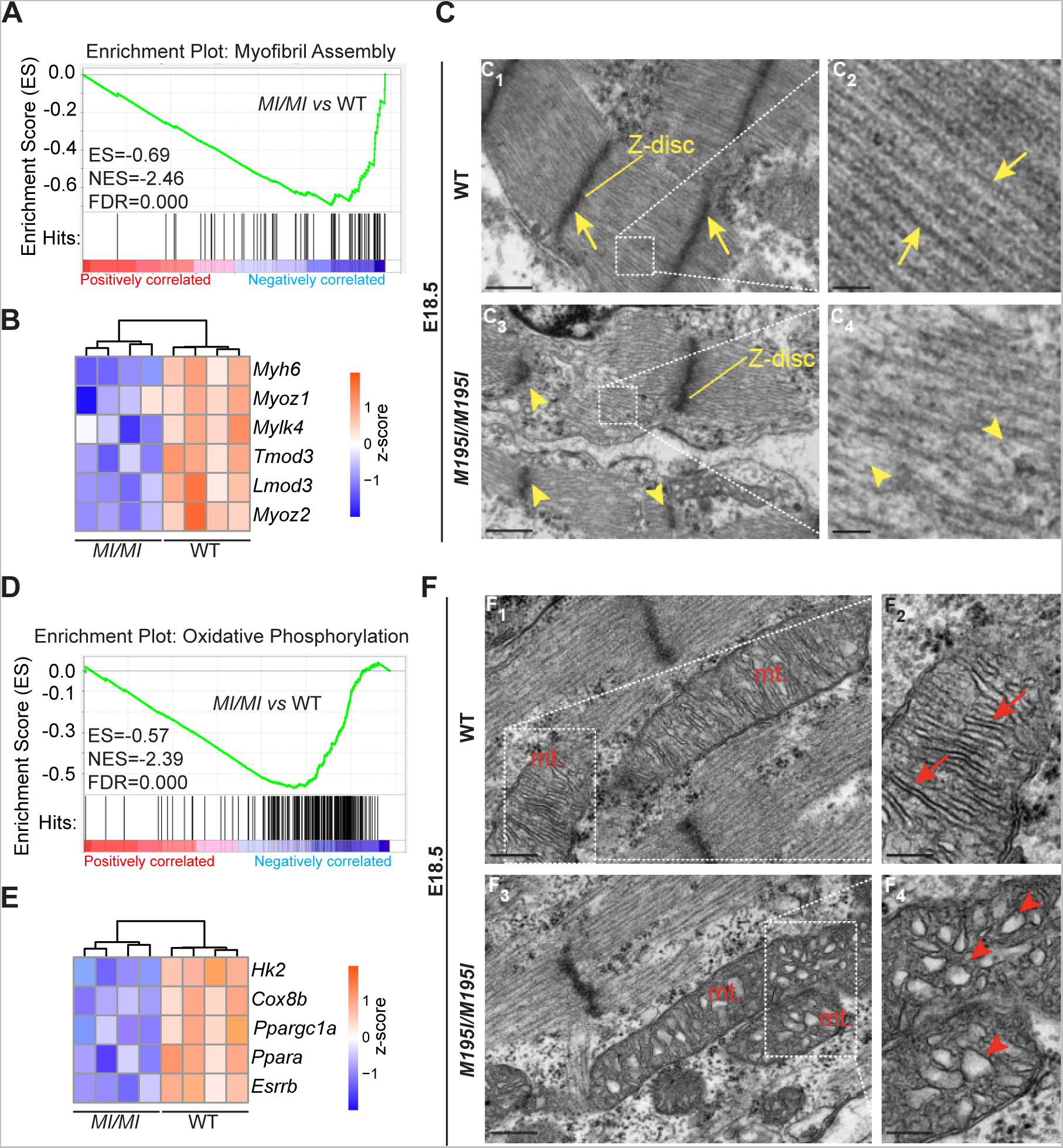
Cardiomyocytes in *CHD4^M195I/M195I^* are immature. **A** and **D**, Gene set enrichment analysis (GSEA) showing the downregulated gene sets of Myofibril Assembly (**A**) and Oxidative Phosphorylation (**D**) in *CHD4^M195I/M195I^*. ES, enrichment score; NES, normalized enrichment score; FDR, false discovery rate. B and E, Heatmaps of representative misregulated genes involved in **A** and **D**. Color bar: z-score. **C**, Representative transmission electron microscopy images of myofibrils in E18.5 hearts. Yellow arrows indicate well-formed Z-disc (**C_1_**) and sarcomeres (**C_2_**) in WT hearts, yellow arrowheads indicate weak, deficient Z-disc formation (**C_3_**) and poorly organized sarcomeres (**C_4_**) in *CHD4^M195I/M195I^* hearts. Scale bars, 500 nm (overview) and 100 nm (magnified). **F**, Representative transmission electron microscopy images of mitochondria (mt.) in E18.5 hearts. Red arrows indicate well-formed mitochondrial cristae (**F_2_**) and red arrowheads indicate poorly organized mitochondrial cristae (**F_4_**). Scale bars, 250 nm (overview) and 100 nm (magnified).

In heart development, cardiomyocyte division is associated with the disassembly of organized contractile fibrils^45, 46^. The observation that *CHD4^M195I/M195I^* cardiomyocytes showed downregulation of gene sets associated with mature cardiomyocytes led us to hypothesize that proliferating cardiomyocytes in *CHD4^M195I/M195I^* remained in an immature state and unable to form functioning sarcomeres. To test this hypothesis, we used transmission electron microscopy to examine the morphology of myofibrils. By E18.5, control hearts were composed of actin filaments assembled into individual contractile units anchored by Z discs (Figure 4C_1_ and 4C_2_). In contrast, *CHD4^M195I/M195I^* hearts displayed a severe reduction in myosin filament density and a concomitant loss or deformation of Z discs (Figure 4C_3_ and 4C_4_). Congruently, in *CHD4^M195I/M195I^* hearts, we found a failure of Tropomyosin to localize into discrete sarcomeres (Figure S9C). These findings suggested that proliferating cardiomyocytes failed to incorporate into organized sarcomeres, which may explain the observation of deficient contractile function in the mutant mice as assessed from echocardiograms (Figure 3).

During maturation, cardiomyocytes undergo metabolic transition from glycolysis to oxidative phosphorylation to produce sufficient ATP to sustain heart function^47, 48^. Consistent with our finding that *CHD4^M195I/M195I^* cardiomyocytes retain properties of immature cells, we found that genes associated with oxidative phosphorylation were underrepresented in the transcriptional profile of *CHD4^M195I/M195I^* hearts (Figure 5D). Prominent examples were downregulation of Hexokinase 2 (*Hk2*) and Cytochrome C Oxidase 8b (*Cox8b*) (Figure 5E), two enzymes essential for metabolism in mature cardiomyocytes^49, 50^. In addition, metabolic transcriptional regulators of mature cardiomyocytes (*Ppargc1a*, *Ppara*, *Esrrb*) were also significantly downregulated in the *CHD4^M195I/M195I^* hearts (Figure 5E). The changes in gene expression in *CHD4^M195I/M195I^* cardiomyocytes correlated with alteration in the maturation of the mitochondria. Mitochondria in WT hearts contained densely organized cristae (Figure 5F_1_ and 5F_2_), the structure housing the electron transport chain and ATP synthase. In contrast, the mitochondrial cristae in the *CHD4^M195I/M195I^* were immature and poorly aligned (Figure 5F_3_ and 5F_4_). Collectively, our molecular and cellular data strongly implied that cardiomyocytes in *CHD4^M195I/M195I^* hearts continued to proliferate until birth and were maintained in an immature state.

### *CHD4^M195I^* leads to a downregulation of ADAMTS and a concomitant ectopic accumulation of the extracellular matrix

A fine-tuned balance between the synthesis and degradation of extracellular matrix (ECM) components is required for proper trabeculation and compaction in developing hearts^13, 51–55^. On the basis of our transcriptional profiling and histological analyses, we hypothesized that the ECM dynamics was dysregulated in *CHD4^M195I/M195I^* hearts. To test this hypothesis, we analyzed the cardiac ECM at E9.5, E10.5, E12.5, and E13.5 with Alcian Blue to detect acidic mucopolysaccharides, including hyaluronan and proteoglycans. *CHD4^M195I/M195I^* hearts showed a robust increase in Alcian Blue in the ventricular cardiac jelly compared with WT at each stage (Figure 6A, Figure S10A, and S10B). We further found an accompanying increase in Versican (Vcan), a large proteoglycan that functions in switching cardiomyocytes from a state of rapid proliferation to differentiation^56^ (Figure 6B-6D, Figure S10A and S10B). Consistently, we found a significant positive correlation between the relative Alcian Blue-positive area and proliferating cardiomyocytes (Ki67^+^/TMY^+^) in *CHD4^M195I/M195I^* hearts (Figure 6E). Thus, cardiomyocyte over-proliferation is associated with ECM accumulation.

**Figure 6.**
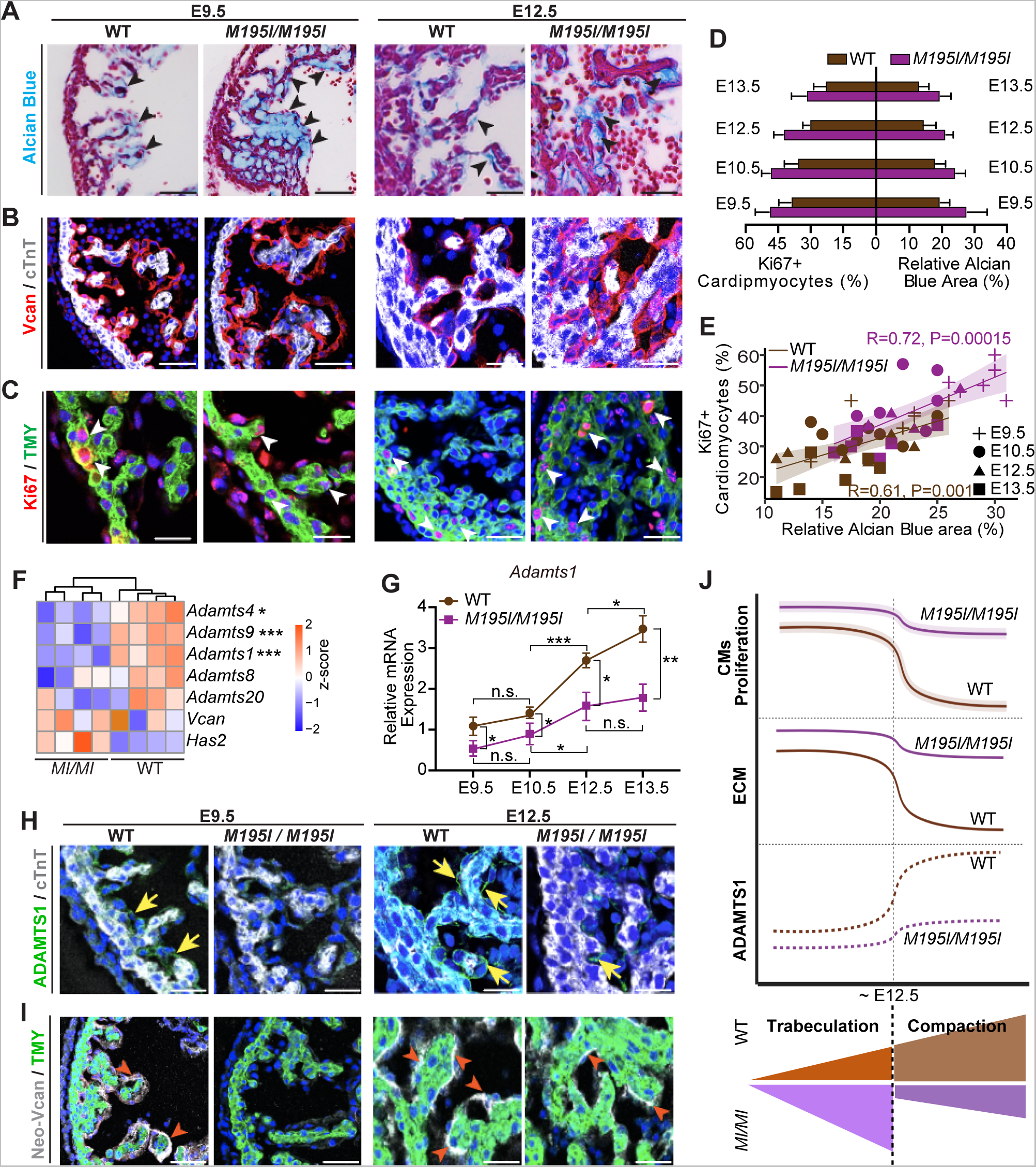
Cardiac ECM dynamics in *CHD4^M195I/M195I^* are dysregulated. **A**, Representative images of Alcian Blue staining for extracellular matrix (ECM) on E9.5 and E12.5 WT and *CHD4^M195I/M195I^* heart paraffin sections. ECM is indicated by black arrowheads. Scale bars, 50 μm (E9.5) and 100 μm (E12.5). **B**, Representative images of immunofluorescent (Versican (Vcan), cardiac Troponin T (cTnT), and DAPI) stained paraffin sections from E9.5 and E12.5 mouse hearts. Scale bars, 50 μm. **C**, Representative images of immunofluorescent-(Ki67, Tropomyosin (TMY), and DAPI) stained paraffin sections from E9.5 and E12.5 mouse hearts. Proliferating cardiomyocytes (Ki67^+^/TMY^+^) are indicated by white arrowheads. Scale bars, 25 μm (E9.5) and 50 μm (E12.5). **D**, Quantification of proliferating cardiomyocyte (Ki67^+^) ratio and relative Alcian Blue-positive area in indicated stages. **E**, Scatter plot showing Ki67^+^ cardiomyocyte ratio relative to Alcian Blue-positive area in WT (brown symbols) or *^M195I/M195I^* (purple symbols) hearts at indicated stages. Pearson correlation coefficients were calculated for determining the correlation. **F**, Heatmap of representative genes identified in E18.5 RNA-seq that are involved ECM dynamics. \**FDR*<0.05, \*\**FDR*<0.01, \*\*\**FDR*<0.001. Color bar: z-score. **G**, RT-qPCR of *Adamts1* expression in WT and *CHD4^M195I/M195I^* hearts at indicated stages (n=3-5 non-pooled hearts per genotype per stage). *Pgk1* gene was used as the internal control, and *Adamts1* expression in E9.5 WT hearts was normalized as 1.0; expression of all other stages and *CHD4^M195I/M195I^* hearts was normalized to E9.5 WT hearts. **H** and **I**, Representative images of immunofluorescent (ADAMTS1, cardiac Troponin T (cTnT), and DAPI, **H**; or Neo-Versican (Neo-Vcan), TMY, and DAPI, **I**) stained sections of E9.5 and E12.5 mouse hearts. ADAMTS1 and Neo-Vcan were indicated by yellow arrows or red arrowheads, respectively. Scale bars, 50 μm. n=6 per genotype per stage in **A** through **E**, **H**, and **I**; data in **D** and **G** are represented as mean ± SEM. Statistical significance was determined with one-way ANOVA (n.s., not significant (P>0.05); \**P*<0.05, \*\**P*<0.01, \*\*\**P*<0.001). J, Proposed schematic for dynamics of cardiomyocyte proliferation, ECM component, ADAMTS1 expression patterns, and sublayers development in WT and *CHD4^M195I/M195I^* hearts.

Interestingly, although we saw a marked increase in Vcan protein in *CHD4^M195I/M195I^* versus WT hearts, we did not observe an increase in Vcan mRNA (Figure 6F and S10C), which suggested that accumulation of Vcan was post-translationally controlled. In parallel, we observed that cardiac expression of ADAMTS1, a critical metalloproteinase for degrading ECM and terminating trabeculation^13, 54^, was significantly lower in the *CHD4^M195I/M195I^* at both gene and protein levels (Figure 6G, H, Figure S10A and S10B). These changes were accompanied by decreased expression of Neoversican (Figure 6I, Figure S10A and S10B), a readout of metalloprotease activity^54^. Taken together, our data suggested that CHD4^M195I^ represses *Adamts1* transcription, at or after E12.5, leading to an accumulation of components of the cardiac jelly (i.e., Vcan), elevated and sustained proliferation of immature cardiomyocytes, and a failure to terminate cardiac trabeculation (Figure 6J).

### Augmented Interaction between CHD4 and BRG1 in *CHD4^M195I/M195I^* hearts

The findings that *Adamts1* was downregulated in *CHD4^M195I/M195I^* cardiomyocytes and the associated increase in the ADAMTS1 substrate Vcan led us to investigate whether *Adamts1* is a direct target of CHD4. The BRG1 component of the multiprotein chromatin remodeling SWI/SNF complexes^57, 58^ represses *Adamts1* in endocardial cells to produce an ECM-rich environment that supports trabeculation. By E12.5, BRG1-mediated repression of *Adamts11* is relieved by an identified mechanism, dissipating the cardiac jelly and preventing excessive trabeculation^54^.

Endogenous BRG1 interacts with CHD4^59, 60^ by directly associating with the N-terminal fragment of CHD4^60, 61^. Coincidently, the CHD4^M195I^ mutation occurs within the CHD4 N-terminus. Thus, we hypothesized that the CHD4^M195I^ protein has a higher affinity than wild-type CHD4 for BRG1, thereby repressing *Adamts1* in *CHD4^M195I/M195I^*.

To test this model, in the presence of Pierce universal nuclease, we isolated the CHD4 cardiac endogenous interactome under physiological conditions from WT and *CHD4^M195I/M195I^* hearts and performed mass spectrometry (MS/MS) analyses. The complexes (N=3) were derived from E13.5 hearts, the time point when *CHD4^M195I/M195I^* cardiomyocytes show an increase in the ADAMTS1 substrate Vcan and have an increase in cardiomyocyte proliferation (Figure 6). We recovered CHD4 from E13.5 cardiac tissue at 57% coverage of the theoretical maximum of 86.6% with trypsin digest of all amino acids (Figure S11). We used an unbiased gene ontology-based bioinformatics classification to screen the functions of proteins associated with CHD4. This analysis showed CHD4 in association with ten components of the SWI/SNF complex, BRG1, SMARCA5, ACTL6A, ARID1A, BAF60C, BAF155, BRD7, BAF57, BAF170, and ARID2. Thus, in E13.5 hearts, CHD4 associated with all the components required for a functional SWI/SNF complex. Moreover, after standardization, *CHD4^M195I/M195I^* was found to be associated with a greater number of spectra and a greater area under the curve for components of the SWI/SNF complex, including BRG1, compared with wild type CHD4 (Figure 7A, Figure S12A and S12B). In aggregate these data indicated that, compared with wild type, CHD4^M195I^ protein may interact with a higher affinity with the SWI/SNF complex in the heart.

**Figure 7.**
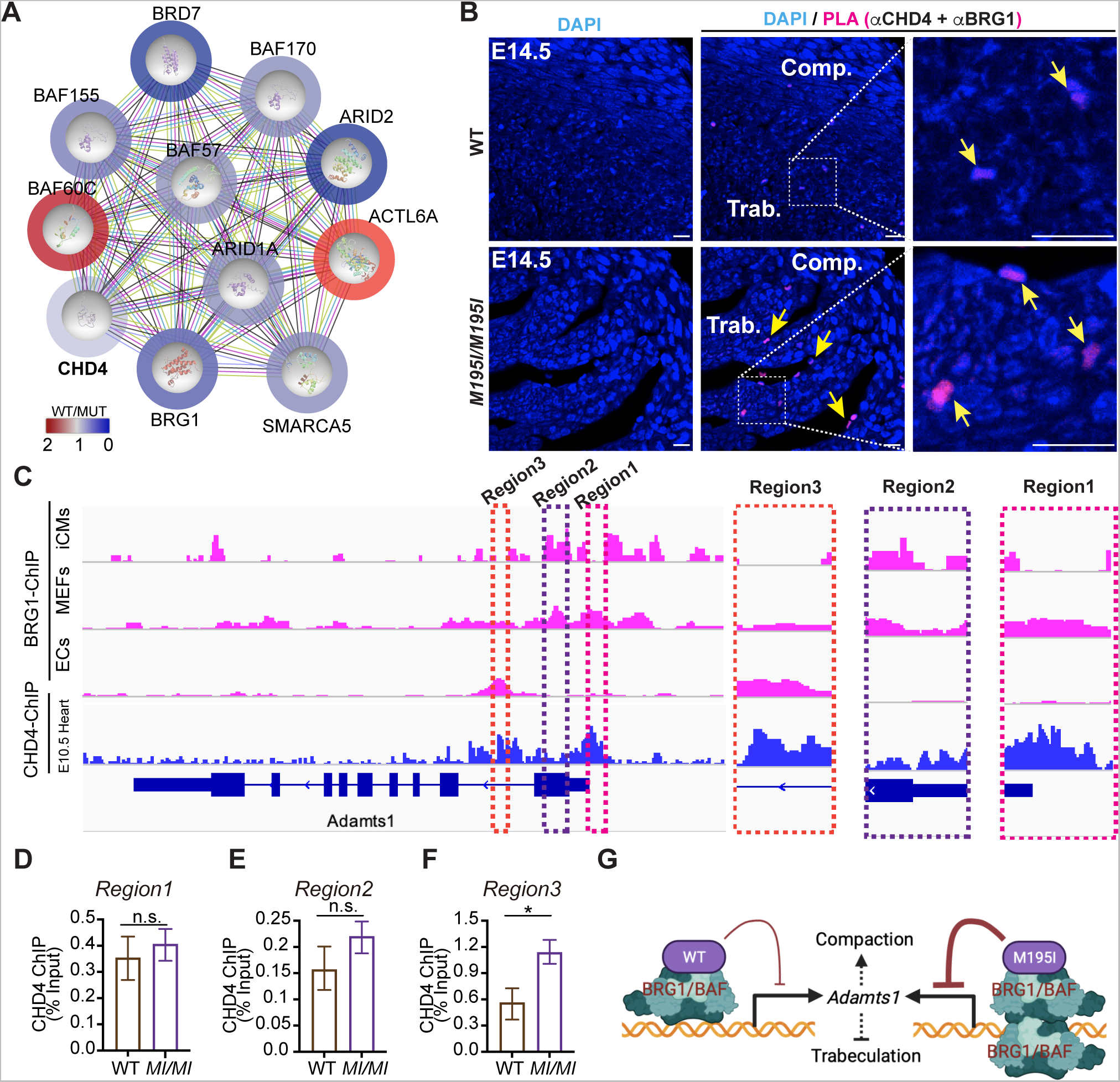
CHD4^M195I^ highly associates with BRG1 to repress *Adamts1*. **A**, Protein network-based prediction with STRING showing CHD4 interactome in E13.5 hearts. Color bar and circles outside of each protein: ratio of M195I CHD4 spectral counts to WT CHD4. Two biological replicates per genotype for the Mass Spectrometry, 15 hearts were pooled for one replicate. **B**, Representative images of in-situ Proximity Ligation Assay (PLA), counterstained with DAPI, performed with anti-CHD4 and anti-BRG1 antibodies on E14.5 WT and *CHD4^M195I/M195I^* heart sections. Scale bars, 20 μm. Positive PLA signals are indicated by yellow arrows; n=3 hearts per genotype. **C**, Visualization by IGV browser of BRG1 or CHD4 ChIP-seq signals across the *Adamts1* locus. Regions of interests are magnified in dashed frames. Data of BRG1 ChIP-seq and CHD4 ChIP-seq were retrieved from published studies: BRG1 ChIP-seq on iCMs (GSE116281)^58^, on MEFs (GSM2671190), on ECs (GSE152892)^81^, CHD4 ChIP-seq on E10.5^26^. **D** through **F**, ChIP-qPCR for CHD4-ChIP samples from E12.5 WT and *CHD4^M195I/M195I^* hearts with primers against the three indicated regions in **C**. Two biological replicates for the ChIP, 15 pooled hearts for each replicate. Data in **D** through **F** are represented as mean ± SEM. Statistical significance was determined with two-tailed unpaired Student’s *t* test (n.s., not significant (P>0.05); \**P*<0.05). **G**, Schematic for regulation of *Adamts1* transcription by CHD4 and BRG1 association.

To confirm the CHD4-BRG1 interaction, we performed an in-situ proximity ligation assay with E14.5 heart sections. We observed interaction between BRG1 and CHD4 in the nuclei at the edge of the trabecular myocardium (Figure 7B) in both WT and mutant hearts (yellow arrows; Figure 7B). Consistent with the IP-MS/MS results, the interaction between BRG1 and mutant CHD4 in the heart was more prominent than the interaction in WT hearts (Figure 7B). These results demonstrated that CHD4 and BRG1 interact in vivo in embryonic heart tissue.

### CHD4^M195I^-BRG1 represses *Adamts1*

Chromatin immunoprecipitation (ChIP)-sequencing reveals that BRG1 and CHD4 co-occupy target genes genome-wide^61, 62^, and that BRG1 and CHD4 dynamically occupy three *cis*-regulatory regions in the *Adamts1* locus^54^, i.e., Regions 1-3 (Figure 7C). To determine whether *CHD4^M195I^* affects binding of CHD4 at *Adamts1*, we performed ChIP-quantitative PCR on WT and *CHD4^M195I/M195I^* hearts at E12.5. We found that *CHD4^M195I/M195I^* had a higher occupancy at Region 3 compared with wild-type CHD4, whereas there was no significant difference at Region 1 or Region 2 (Figure 7D-7F). These results suggested that *CHD4^M195I/M195I^* associates more frequently or with a greater affinity with BRG1 at the Region 3 locus, thereby maintaining repression of *Adamts1* (Figure 7G).

### Restoration of ADAMTS1 Rescues Overgrowth of Trabeculae

Our data suggested that ADAMTS1 has an essential function in dissipating the cardiac jelly and preventing excessive trabeculation^54^. To test this hypothesis in vivo, we examined the ability of recombinant ADAMTS1 (rADAMTS1) to restore the termination of trabeculation in hearts derived from *CHD4^M195I/M195I^* E11.5 embryos. For these studies, we first used heart explant cultures^63–67^ in which the right ventricles were isolated and cultured in the presence of bead grafts that contained rADAMTS1 or BSA (Figure 8A). We observed that rADAMTS1 significantly inhibited the expression of Vcan and reduced the number of cycling cardiomyocytes (Figure 8B_1_, 8B_2_ and 8C). Importantly, the changes in Vcan expression were not associated with changes in the proliferation of epicardial cells (Figure 8D), which suggested that rADAMTS1 acts in a cardiac cell type-specific manner. Thus, these data demonstrated that restoration of ADAMTS1 to the endocardial environment of *CHD4^M195I/M195I^* hearts inhibited ECM accumulation and cardiomyocyte proliferation (Figure 8B_3_ and 8B_4_).

**Figure 8.**
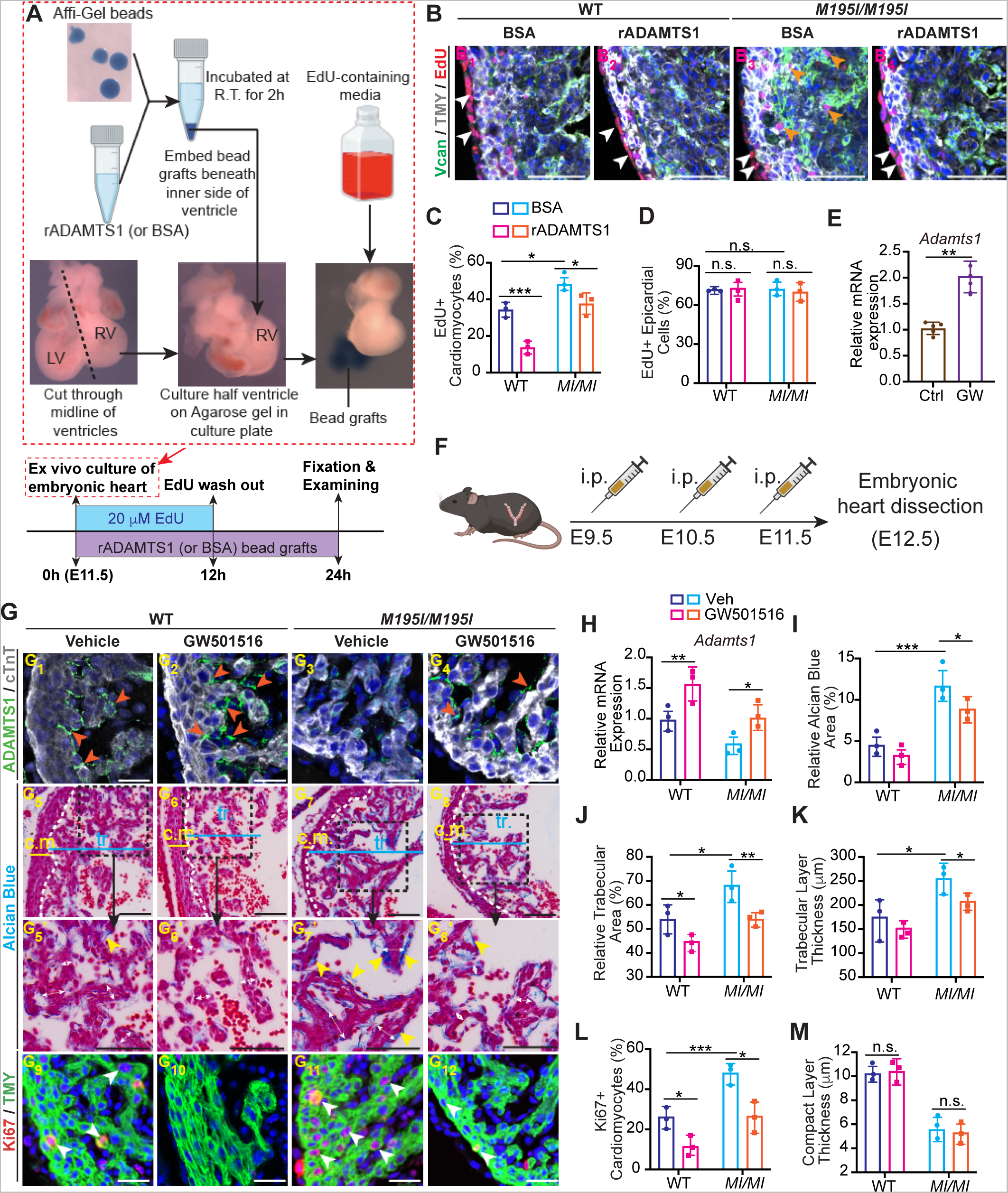
Restoration of ADAMTS1 rescues hypertrabeculation in *CHD4^M195I/M195I^*. **A**, Workflow for treating the explant embryonic heart ventricles with recombinant ADAMTS1 protein. Hearts were dissected at E11.5 and cut through the middle line of the ventricles; the right ventricles were cultured on low-melting point agarose gel. Recombinant ADAMTS1 protein (BSA was used as control) was incubated with the Affi-Gel bead at ambient temperature for 2 hours, and then the ADAMTS1 bead grafts were embedded beneath the inner side of the cultured ventricles for 24 hours. Explant ventricles were cultured in EdU-containing media for the first 12 hours, and then cultured with EdU-free media for another 12 hours. **B**, Representative images of immunofluorescent-(EdU, Vcan, TMY, and DAPI) stained sections of ventricle explants. EdU^+^ cardiomyocytes and epicardial cells are indicated by orange and white arrowheads, respectively. Scale bars, 50 μm. **C** and **D**, Quantification of EdU^+^ cardiomyocytes (**C**) and epicardial cells (**D**) in the explants of four conditions in **B** (n=3 per condition). **E**, RT-qPCR of *Adamts1* expression in ADAMTS1 agonist (GW501516) or vehicle treated WT heart explants. *Pgk1* gene was used as the internal control; n=5 explants per group. **F**, Workflow for treating pregnant *CHD4^M195I/+^* dams with GW501516. *CHD4^M195I/+^* mice were intercrossed for timed-mating, and GW501516 was administered to the plugged *CHD4^M195I/+^* mice by intraperitoneal injection at E9.5, E10.5, and E11.5. Embryonic hearts were dissected and examined at E12.5 (n=4 WT and 4 *CHD4^M195I/M195I^* embryonic hearts were examined from three GW501516-treated *CHD4^M195I/+^* dams). **G**, Representative images of ADAMTS1, cTnT immunofluorescence (**G_1_** through **G_4_**), Alcian Blue staining (**G_5_** through **G_8_’**), and Ki67, TMY immunofluorescence (**G_9_** through **G_12_**) performed on embryonic hearts that were dissected from GW501516-treated *CHD4^M195I/+^* dams. ADAMTS1^+^ stains are indicated by orange arrowheads (**G_1_** through **G_4_**). Thickness of trabeculae are indicated by white double-headed arrows, and Alcian Blue-positive stains are indicated by yellow arrowheads (**G_5_** through **G_8_’**). EdU^+^ cardiomyocytes are indicated by white arrowheads (**G_9_** through **G_12_**). Scale bars, 50 μm (**G_1_** through **G_4_**, **G_9_** through **G_12_**) and 100 μm (**G_5_** through **G_8_’**). **H**, RT-qPCR of *Adamts1* expression in the embryonic hearts of four conditions. *Pgk1* gene was used as the internal control. **I** through **M**, Quantification of relative Alcian Blue area (**I**), relative trabecular area (**J**), trabeculae thickness (**K**), Ki67^+^ cardiomyocyte ratio (**L**), and compact layer thickness (**M**). n=3 independent hearts for each condition in **G** through **M**. All data in **C** through **E**, **H** through **M**, are represented as mean ± SEM. Statistical significance in **C**, **D**, **H** through **M**, was determined with two-way ANOVA, in **E** was determined with two-tailed unpaired Student’s *t* test (n.s., not significant (P>0.05); \**P*<0.05, \*\**P*<0.01, \*\*\**P*<0.001).

Studies with breast cancer cell lines demonstrated that the pharmacological compound GW501516 acts by transcriptionally upregulating ADAMTS1^68^. We tested the ability of the GW501516 to upregulate ADAMTS1 in heart tissue. RT-PCR of *Adamts1* derived from control and GW501516 treated heart explants confirmed that GW501516 significantly upregulated *Adamts1* mRNA in cardiac tissue (Figure 8E). Next, we tested the ability of GW501516 to restore cessation of trabeculation in utero. To this end, we intercrossed *CHD4^M195I/+^* mice and injected GW501516 intraperitoneally into pregnant *CHD4^M195I/+^* females carrying litters between stages E9.5 to E11.5 (once a day; Figure 8F). At E12.5, we found that ADAMTS1 mRNA and protein were significantly induced in cardiomyocytes by GW501516 in the WT and *CHD4^M195I/M195I^* hearts (Figure 8G_1_-8G_4_ and 8H). Strikingly, cardiomyocyte proliferation decreased and there was a concomitant decrease in the thickness of trabeculae layer upon GW501516 treatment of *CHD4^M195I/M195I^* hearts (Figure 8G_5_-8G_12_, 8I-8L). However, GW501516 treatment did not affect the thickness of compact myocardium (Figure 8M). These studies indicated that ADAMTS1 regulates the mid-gestation cessation of trabeculae growth. These data further indicated that administration of ADAMTS1 in culture or in utero led to an amelioration of key aspects of LVNC, partly by regulation in *CHD4^M195I^* hearts.

## Discussion

Mutations in chromodomain helicase DNA-binding protein 4 (CHD4), the catalytic component of nucleosome remodeling deacetylase (NuRD), leads to congenital heart disease, including atrial and ventricular septal defects^26, 28, 32, 69, 70^. However, CHD4 is expressed in most cell types; thus, it was not known how patients with Chd4 missense mutations display a restricted cardiac phenotype. In this study, we identified a proband for CHD4 (CHD4^M202I^) who had cardiac abnormalities, and we generated a mouse patient-specific model for this mutation (CHD4^M195I^). Using our patient model, we established the mechanisms by which CHD4^M195I^ leads to impaired cardiac function, and we identified potential approaches to LVNC therapy. Critically, we showed that administration of ADAMTS1 in culture or in utero rescued aspects of the LVNC associated phenotype.

### Left ventricular noncompaction

Whether and how cardiomyocyte proliferation leads to LVNC hypertrabeculation are controversial issues. Recent fate mapping studies revealed that abolishing proliferation in the compact layer can lead to a hypertrabeculation phenotype in mouse^71^. Consistently, cardiac conditional mutations in *Jag1*, *Jag2*, *Prdm16, and RBPMS* all lead to a decrease in proliferation and hypertrabeculated hearts^10, 14, 72^. Conversely, other studies suggest that trabeculation occurs at the expense of compact layer. For example, Luxan et al. demonstrated that mutations in the E3 ubiquitin ligase MIB1 led to LVNC in mouse and human. Enlarged noncompacted trabeculae in MIB1 mice showed an increase in cardiomyocyte proliferation and a downregulation of Notch activity ^15^. Analogously, deletion of *Nkx2-5* in trabecular myocardium^73^ and global knockout of *Fkbp1a*^74^ or *Plxnd1*^53^ all led to hypertrabeculation and reduced compaction that is associated with an increase in cardiomyocyte proliferation. One potential contradiction in these latter findings is that there was an increase in proliferation of cardiomyocytes in the compact layer, yet, the compact layer was thinner than in controls. We favor the explanation that cardiomyocyte proliferation leads to noncompaction akin to a zebrafish model in which proliferation-induced cellular crowding at the tissue scale triggers tension heterogeneity among cardiomyocytes. The crowding in turn drives cardiomyocytes with higher contractility to delaminate and seed the trabecular layer^75^. Thus, *CHD4^M195I^* leads to proliferation and heterogenicity of cardiomyocytes, which leads to a greater number of cardiomyocytes that enter and proliferate within the trabecular layer.

### ADMATS1 and LVNC

ADMATS1 is a member of the ADAMTS (a disintegrin and metalloproteinase with thrombospondin motif) protein family^76^. Here we demonstrated that *CHD4^M195I/M195I^* led to a downregulation of ADAMTS1 and an accumulation of ECM components. Furthermore, administration of ADAMTS1 attenuated key LVNC properties of *CHD4^M195I/M195I^* hearts.

Disruption of Notch signaling in myocardium or endocardium leads to cardiac noncompaction^14, 15, 53, 74, 77–80^. In the present study, transcriptional profiling of *CHD4^M195I/M195I^* hearts revealed only a modest decrease in a limited number of components of the Notch pathway, e.g., *Dll4* and *Jag2*. The *CHD4^M195I^* mutation occurs in a highly conserved N-terminal region of human CHD4 termed CHD4-N (residues 145–225). In vitro, CHD4-N binds poly (ADP-ribose) with higher affinity than it binds DNA, suggesting that CHD4-N recognizes the DNA backbone instead of making specific interactions with the nucleotides. Whether the M202I mutation affects CHD4’s affinity or recruitment to DNA remains to be determined. Our findings suggest that *CHD4^M195I^* acts either downstream or in parallel to Notch signaling. Thus, it will be interesting to determine whether administration of ADAMTS1 rescues LVNC defects that result from mutations in Notch pathway components.

### Conclusions

Our *CHD4^M195I/M195I^* mouse model recapitulated the ventricular noncompaction abnormality in human LVNC. Thus, *CHD4^M195I/M195I^* provided comprehensive understanding of the mechanisms of hypertrabeculation and ventricular noncompaction, and the model defined a previously unknown function of CHD4 in heart development and diseases. Moreover, the observation that administration of ADAMTS1 ameliorated aspects of LVNC suggests a possible ADAMTS1-based therapeutic approach for LVNC patients.

## Nonstandard Abbreviations and Acronyms

ADAMTS1: ADAM metallopeptidase with thrombospondin type 1 motif 1
BRG1: SWI/SNF related, matrix associated, actin dependent regulator of chromatin, subfamily A, member 4
CHD4: chromodomain helicase DNA binding protein 4
ChIP: chromatin immunoprecipitation
cTnT: cardiac troponin T
ECM: extracellular matrix
EdU: 5-ethynyl-2’-deoxyuridine
E.F.: ejection fraction
F.S.: fractional shortening
GSEA: gene set enrichment analysis
IVS: interventricular septum
IP/MS: immuno-purification/mass spectrometer
LA: left atrium
LV: left ventricle
LVNC: left ventricular noncompaction
NuRD: nucleosome remodeling deacetylation
RA: right atrium
RV: right ventricle
TMY: tropomyosin
VCAN: versican
WGA: wheat germ agglutinin

## Acknowledgments

We greatly thank the generation of the *CHD4^M195I^* mouse line by the U.N.C. Animal Model Core. We thank the U.N.C. Microscopy Services Laboratory Core for the processing and imaging of the transmission electron microscopy, and we also thank the performance of proteomics analysis by the U.N.C. Michael Hooker Proteomics Center.

## Supplemental Material

### Supplemental Figures

**Figure S1.**
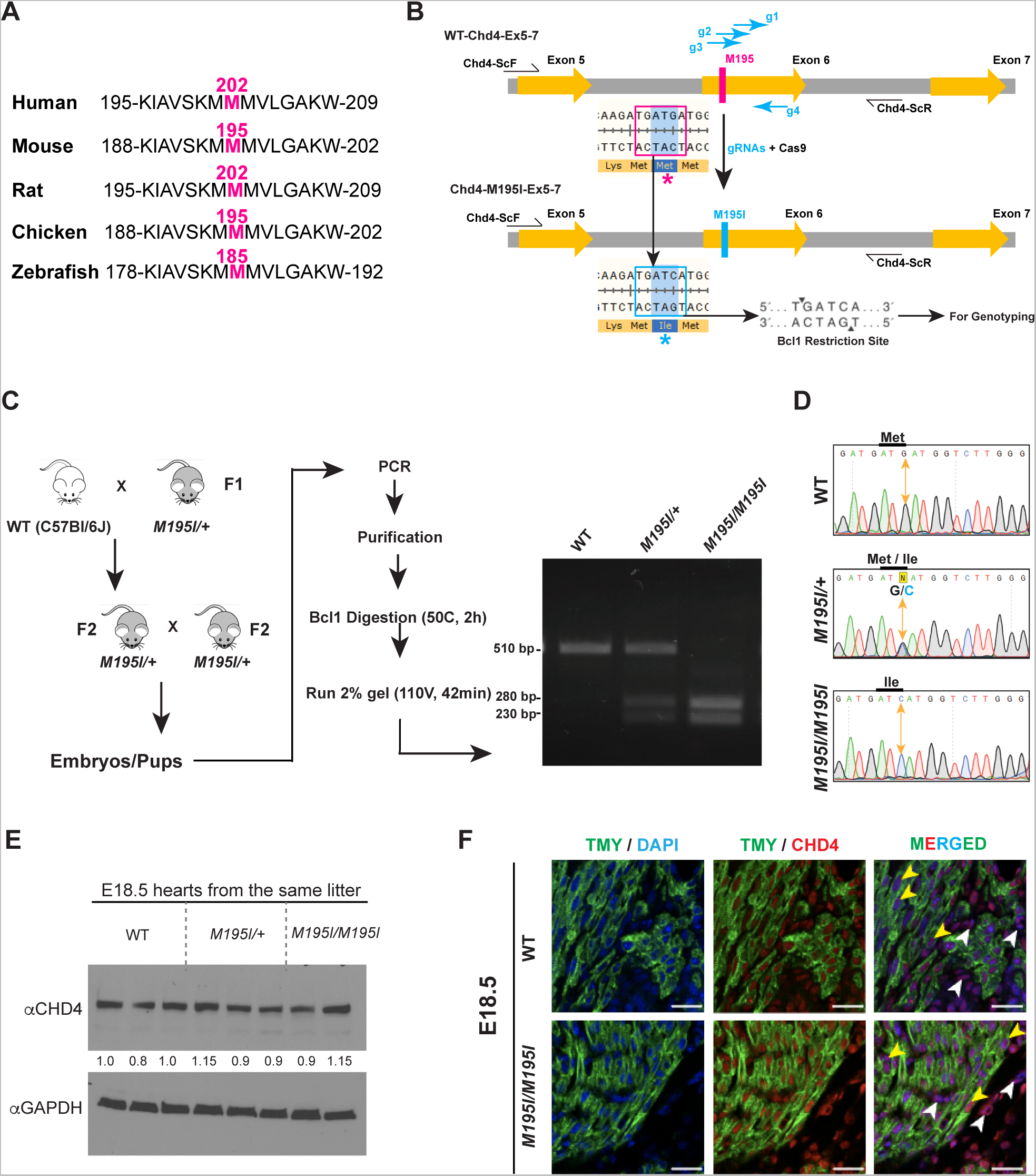
Generation and validation of *CHD4^M195I^* mutant mouse line. **A**, Protein sequence alignment of CHD4 orthologs across indicated species. **B**, Schematic of generating *CHD4^M195I^* mouse line with CRISPR/CAS9 technology. **C**, Mouse breeding strategy and genotyping workflow. **D**, Validation of each genotype by sequencing. **E**, Western blotting performed on hearts of littermates at E18.5 showing no significant differences in CHD4 protein expression. Bands intensity was quantified with Image J and normalized to GAPDH. **F**, Representative images of CHD4 and Tropomyosin (TMY) immunofluorescence performed on E18.5 WT and *CHD4^M195I/M195I^* heart sections, yellow arrowheads indicate cardiomyocytes (TMY^+^), white arrowheads indicate noncardiomyocytes (TMY^-^). Scale bars, 25 μm.

**Figure S2.**
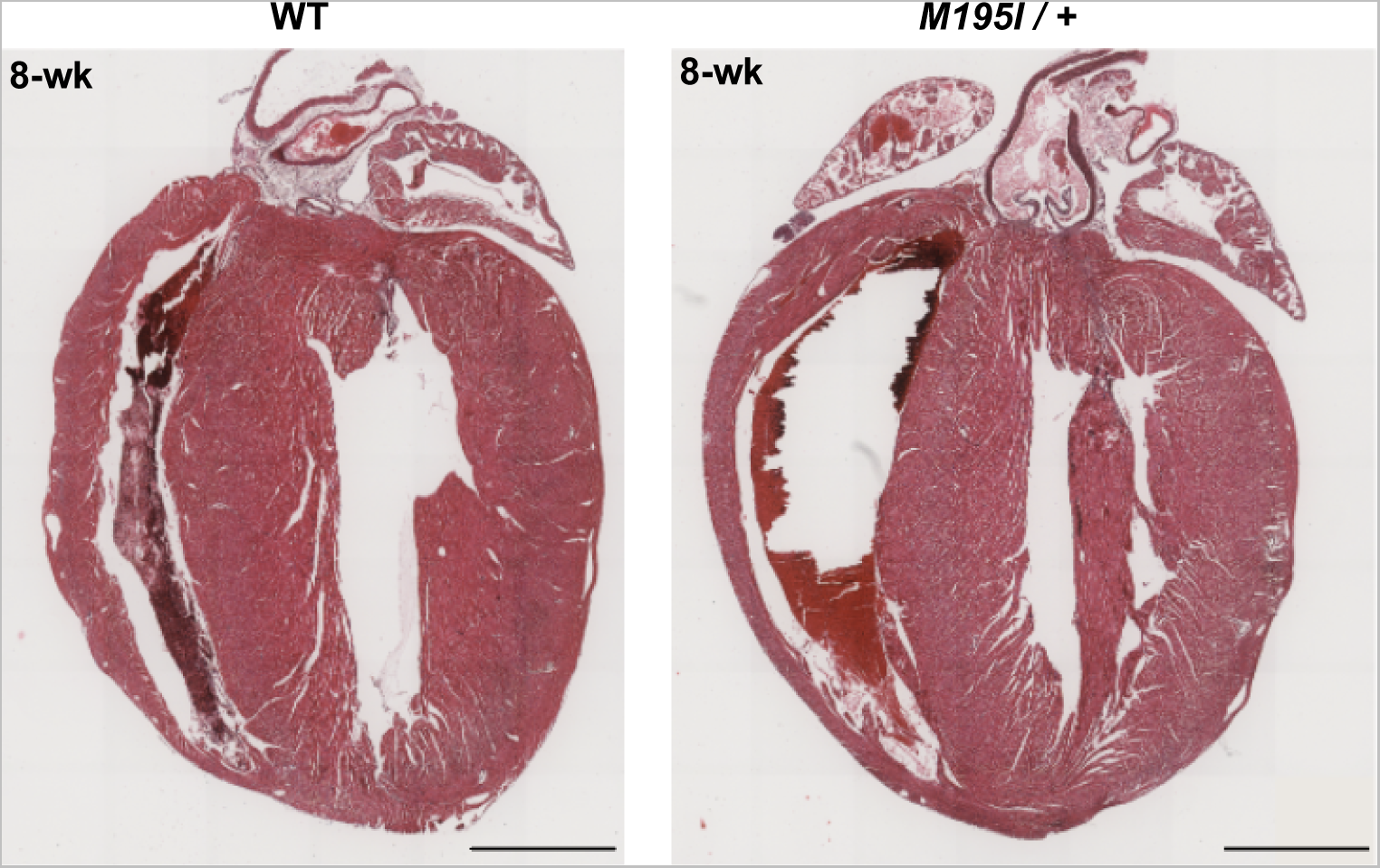
Histological analysis on 8-week hearts. Representative H&E-stained images of WT and *CHD4^M195I/+^* heart sections at 8 weeks. RA, right atrium; LA, left atrium; RV, right ventricle; LV, left ventricle; IVS, interventricular septum. Scale bars, 2mm.

**Figure S3.**
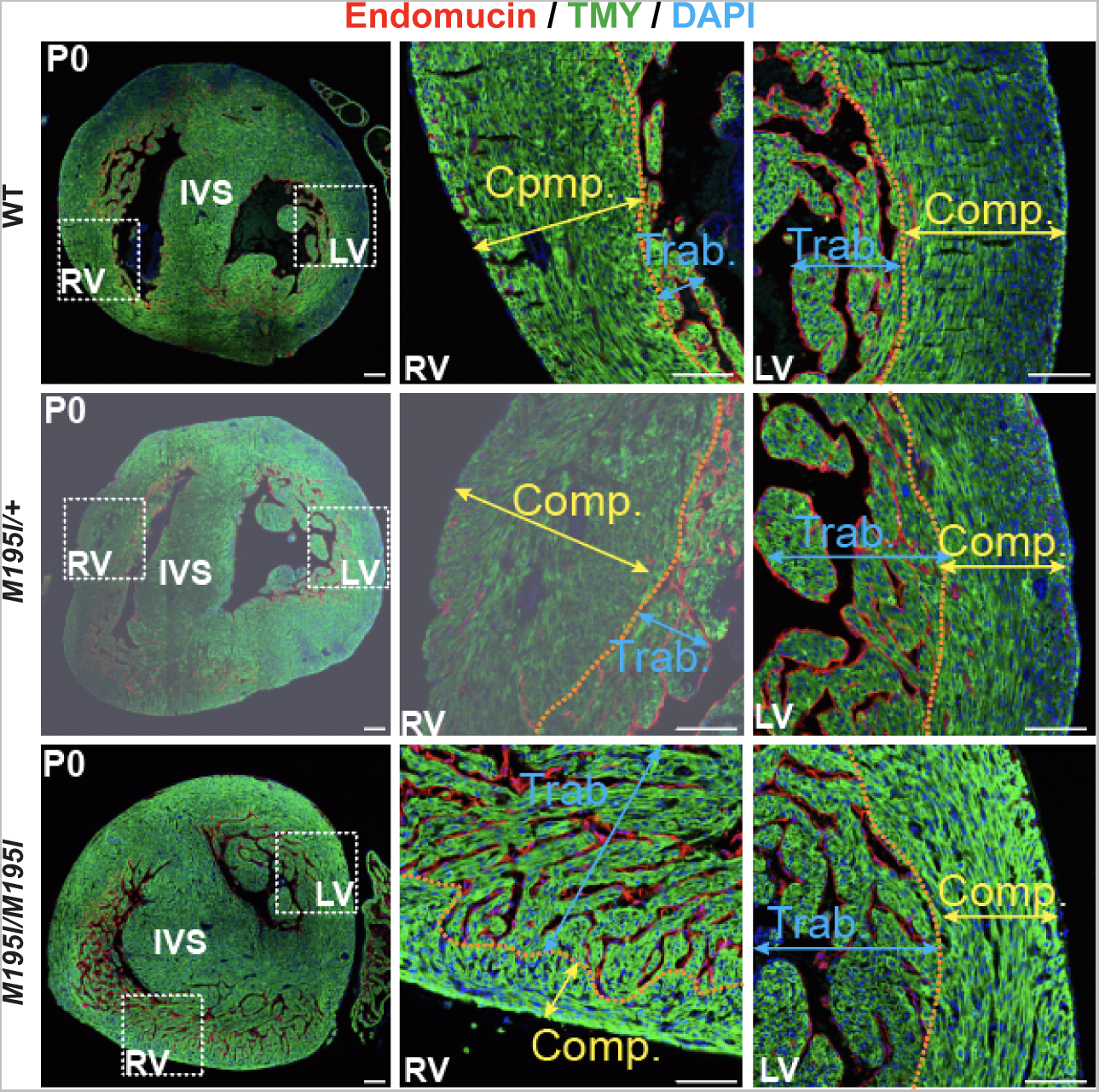
Immunofluorescent analysis on transverse sections of P0 hearts. Representative immunofluorescent (Endomucin, TMY, and DAPI) stained transverse paraffin sections from P0 WT, *CHD4^M195I/+^*, and *CHD4^M195I/M195I^* hearts (n=3 independent hearts per genotype). The approximal boundaries between compact myocardium (Comp.; indicated by yellow double-headed arrows) and trabecular myocardium (Trab.; indicated by blue double-headed arrows) are indicated by orange dashed lines. RV, right ventricle; LV, left ventricle; IVS, interventricular septum. Scale bars: 100 μm.

**Figure S4.**
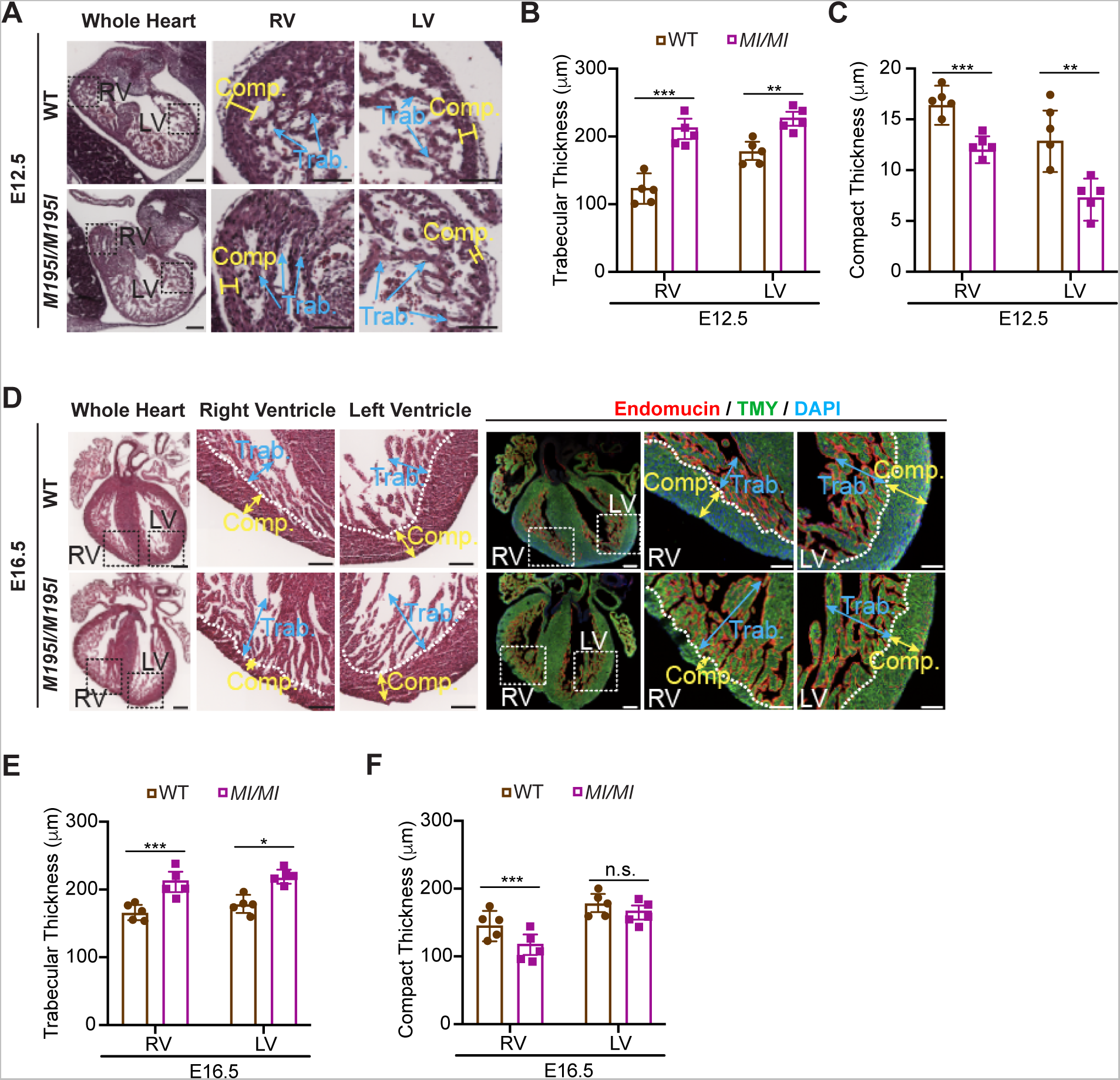
Ventricular noncompaction of *CHD4^M195I/M195I^* hearts at E12.5 and E16.5. **A**, Representative images of H&E-stained paraffin sections of WT and *CHD4^M195I/M195I^* hearts at E12.5, scale bars, 50 μm. **D**, Representative images of H&E-stained and immunofluorescent (Endomucin, TMY, and DAPI) stained paraffin sections of WT and *CHD4^M195I/M195I^* hearts at E16.5, the approximal boundaries between compact myocardium (Comp.; indicated by yellow double-headed arrows) and trabecular myocardium (Trab.; indicated by blue double-headed arrows) are indicated by white dashed lines. Scale bars, 100 μm. **B**, **C**, **E**, and **F**, Measurements of sublayer thicknesses of WT and *CHD4^M195I/M195I^* heart sections from E12.5 (**B** and **C**) and E16.5 (**E** and **F**). Data in **B**, **C**, **E**, and **F** are represented as mean ± SEM, n=5 for each genotype per stage. Statistical significance was determined with two-tailed Student’s *t* test (n.s., not significant (P>0.05); *P<0.05, **P<0.01, ***P<0.001).

**Figure S5.**
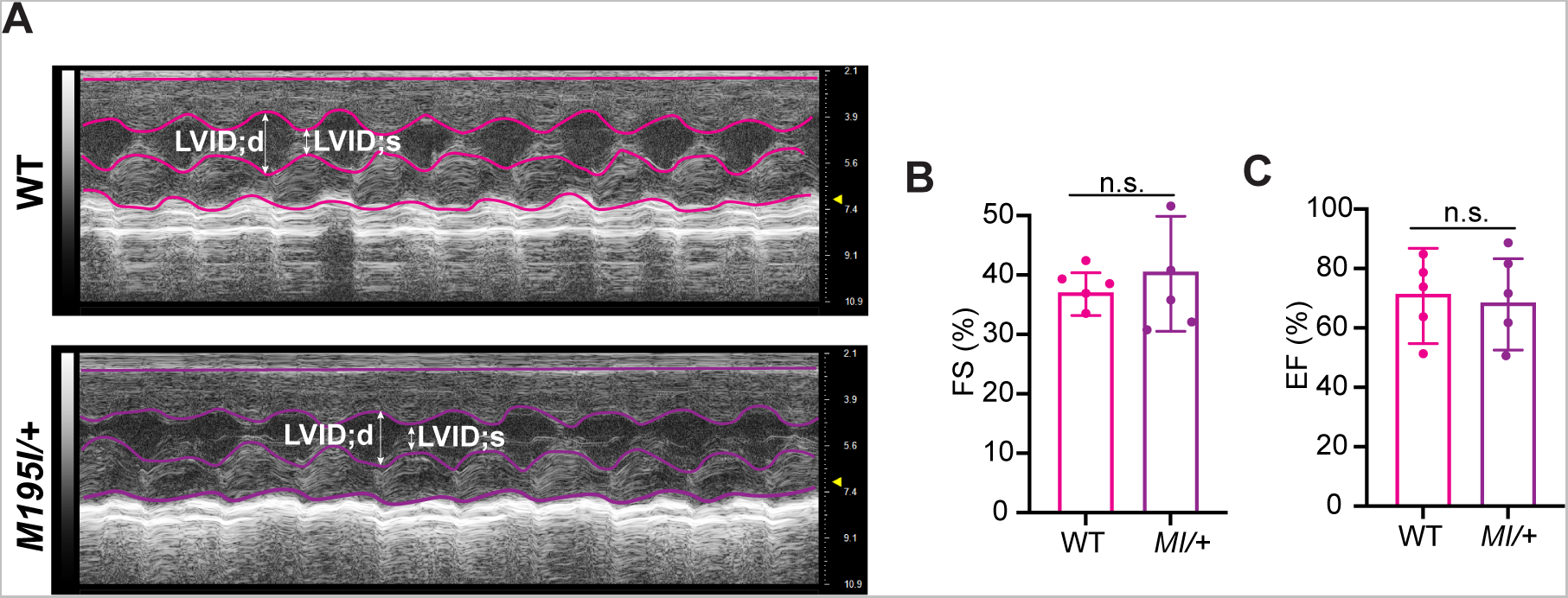
No significant abnormalities in cardiac functions between adult WT and *CHD4^M195I/+^* hearts. **A**, Representative examples of M-mode echocardiography of WT and *CHD4^M195I/+^* mice at 8 weeks. LVID;d: end-diastolic left ventricular internal diameter; LVID;s: end-systolic left ventricular internal diameter. **B** and **C**, Quantification of left ventricular fractional shortening (FS) (**B**), left ventricular ejection fraction (EF) (**C**) for the examined mice (n=5 independent mice per genotype). Data are represented as mean ± SEM. Statistical significance was determined with unpaired Student’s *t* test (n.s., not significant(P>0.05)). *MI/+*: *CHD4^M195I/+^*.

**Figure S6.**
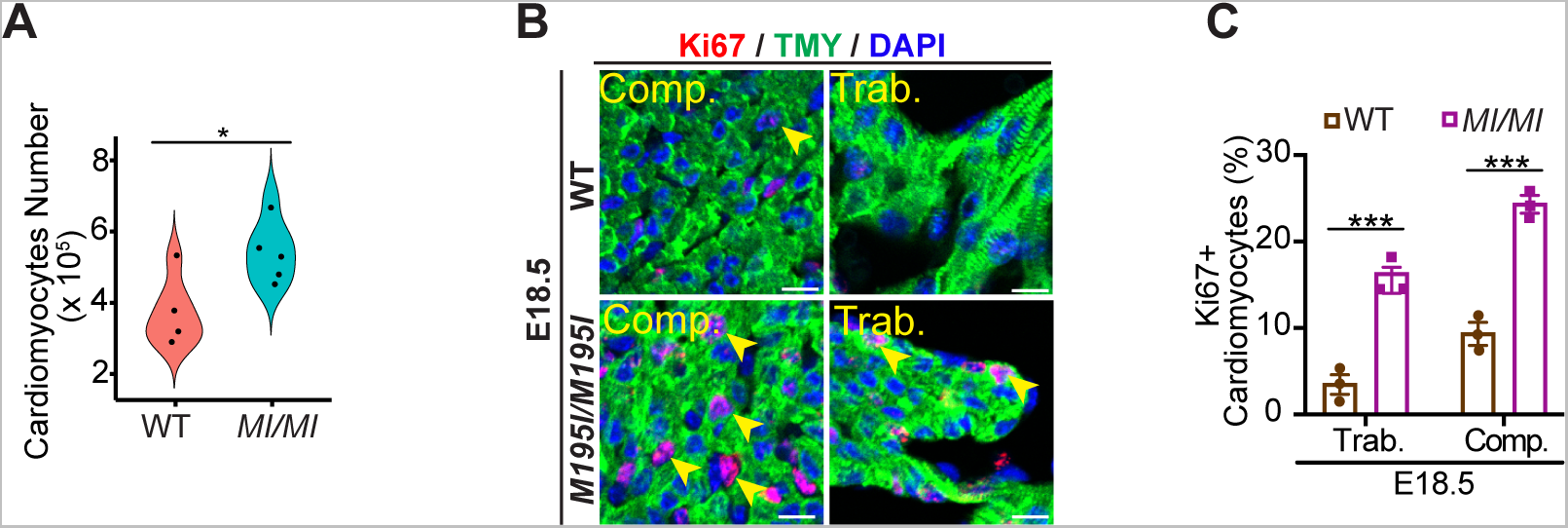
Increased cardiomyocyte proliferation in *CHD4^M195I/M195I^* heart. **A**, Violin plot showing numbers of isolated cardiomyocytes from E18.5 WT and *CHD4^M195I/M195I^* hearts. n=4 independent hearts for WT, n=5 independent hearts for *CHD4^M195I/M195I^*. **B**, Representative images of immunofluorescent (TMY, Ki67, and DAPI) stained paraffin sections from E18.5 WT and *CHD4^M195I/M195I^* mouse hearts. Proliferating cardiomyocytes were indicated by yellow arrowheads. Scale bars, 20 μm. **C**, Quantification of proliferating cardiomyocytes ratio (Ki67^+^/total cardiomyocytes) in E18.5 WT and *CHD4^M195I/M195I^* hearts (n=3 per genotype). Data in **A** and **C** are represented as mean ± SEM, evaluated by two-tailed unpaired Student’s *t* test (*P<0.05, ***P<0.001). *MI/MI*: *CHD4^M195I/M195I^,* Trab.: trabecular, Comp.: compact.

**Figure S7.**
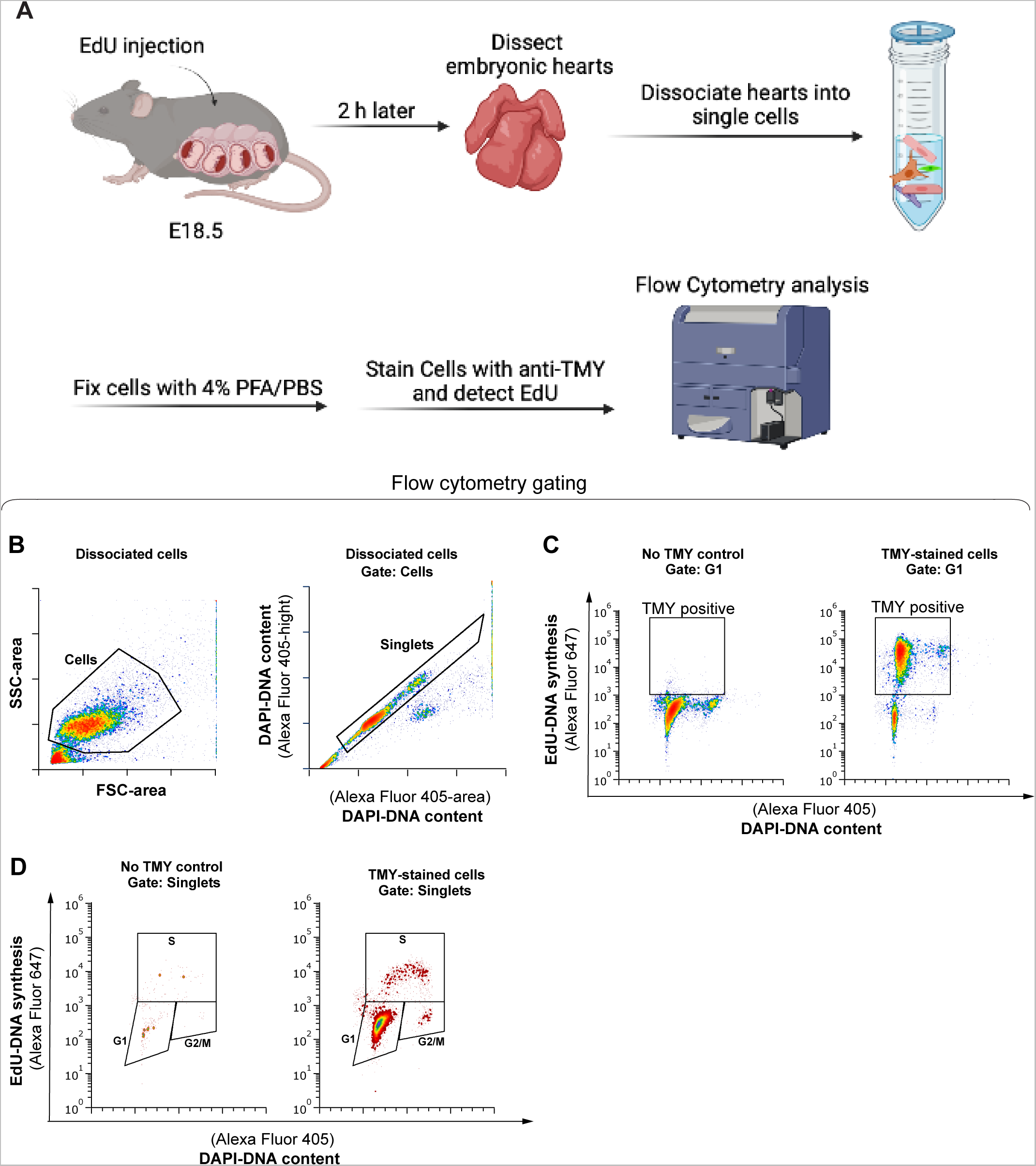
Cell cycle profiling for cardiomyocytes derived from *CHD4^M195I/M195I^* and WT hearts. **A**, Workflow for the cell cycle profiling assay. Schematics was Made with *Biorender*. **B**, Gating to discriminate cells from debris was FS-area vs SS-area (left), singlets (individual cells) from clumps of cells/doublets using DAPI height vs DAPI area (right). **C**, Non-specific background staining by the secondary antibody was measured using a negative control sample stained without primary abtibody, TMY (left) or with TMY (right). Only TMY+ cells were analyzed for cell cycle phases in panel **D**. **D**, Cell cycle phases were determined with DAPI and EdU. Negative control to define background staining without TMY (left). Cells were stained with both TMY and EdU were analyzed for cell cycle phases distribution (right). Background thresholds were set using these controls and applied to all experimental samples.

**Figure S8.**
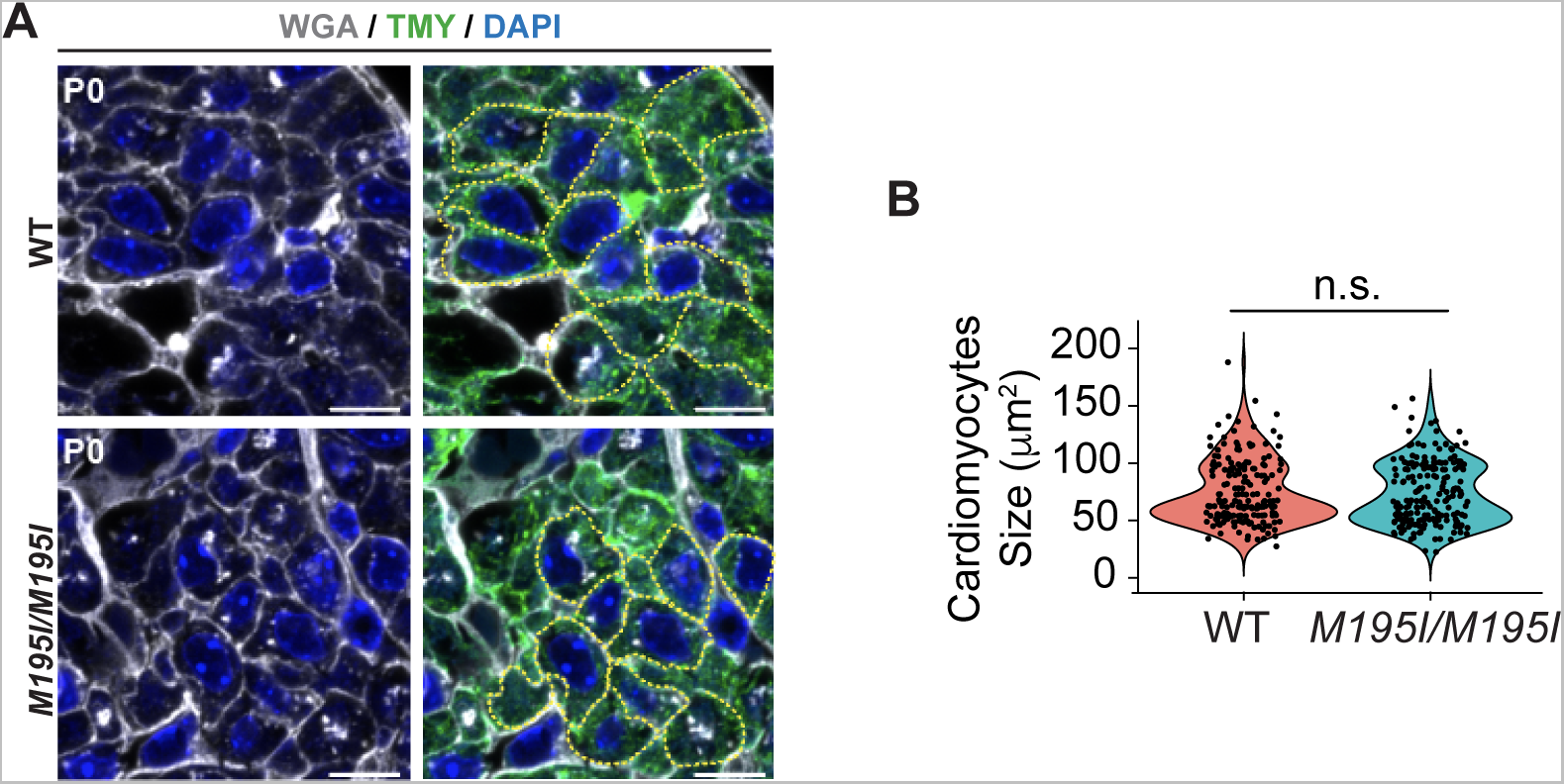
No changes in cardiomyocyte size in *CHD4^M195I/M195I^* heart. **A**, Representative images of immunofluorescent (Wheat Germ Agglutinin (WGA), TMY, and DAPI) stained paraffin transverse sections of WT and *CHD4^M195I/M195I^* hearts at P0. Representative cardiomyocytes are outlined with yellow dashed curve aligning WGA stains. Scale bars, 10 μm. **B**, Quantification of cardiomyocyte size in the P0 WT and *CHD4^M195I/M195I^* mouse heart transverse sections (n=100 cardiomyocytes were measured from 6 hearts per genotype). Statistics was evaluated by two-tailed unpaired Student’s *t* test (n.s., not significant (P>0.05)).

**Figure S9.**
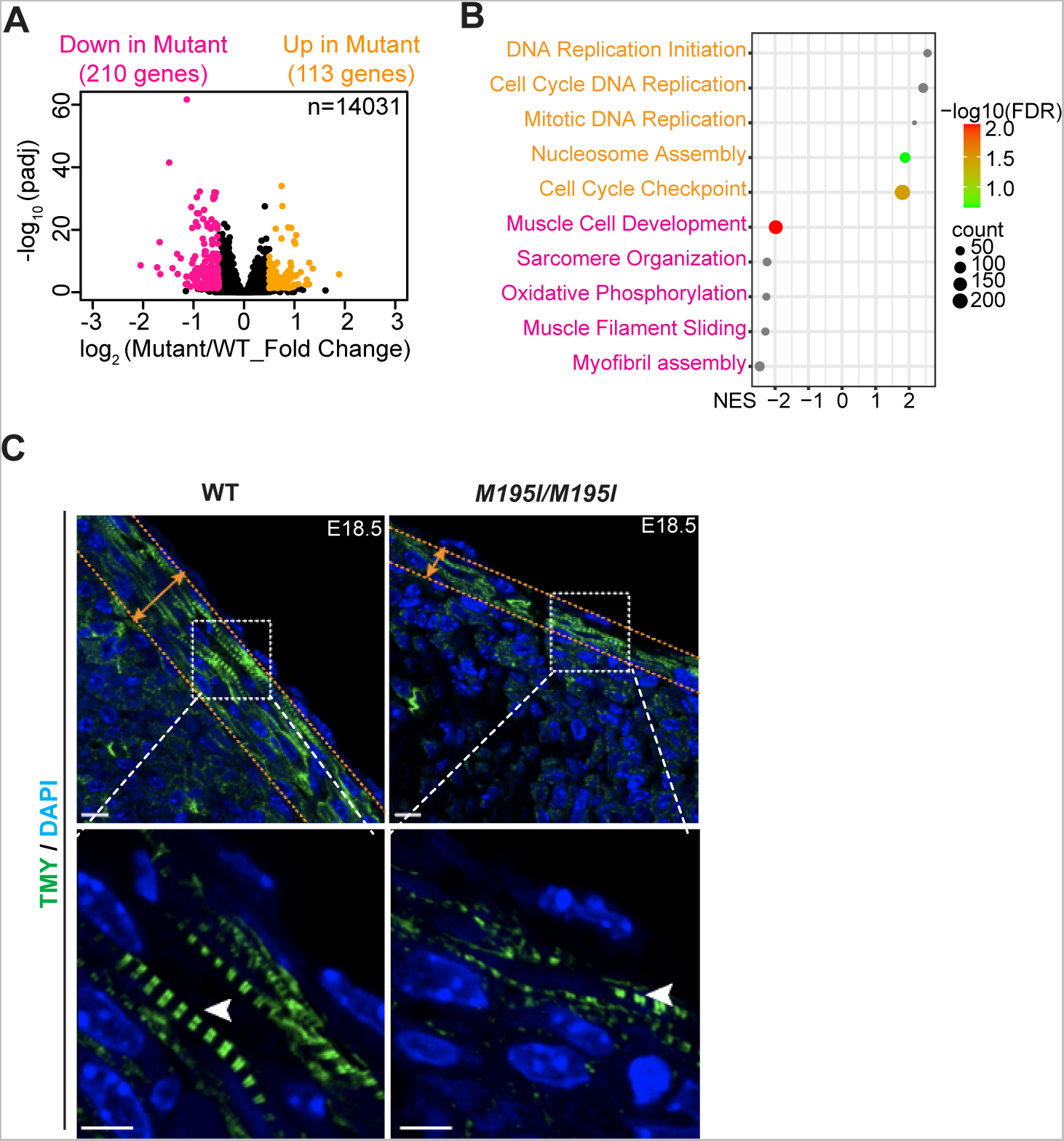
RNA-seq analysis in WT and *CHD4^M195I/M195I^* hearts. **A**, Volcano plot for genes identified in RNA-seq at E18.5. Genes significantly downregulated [adjusted P value <0.05, log_2_ (Fold Change) < −0.5] in the *CHD4^M195I/M195I^* hearts are in pink, genes significantly upregulated [adjusted P value <0.05, log_2_ (Fold Change) > 0.5] in *CHD4^M195I/M195I^* hearts are yellow. **B**, Enriched gene sets obtained from GESA analysis. Upregulated pathways in *CHD4^M195I/M195I^* hearts are in yellow, downregulated pathways in *CHD4^M195I/M195I^* hearts are in pink. NES, normalized enrichment score. **C**, Representative immunofluorescent (TMY and DAPI) stained images of E18.5 WT and *CHD4^M195I/M195I^* heart sections. The contractile apparatus are outlined by orange dashed lines and double-headed arrows. Sarcomeres are indicated by white arrowheads. Scale bars, 5 μm.

**Figure S10.**
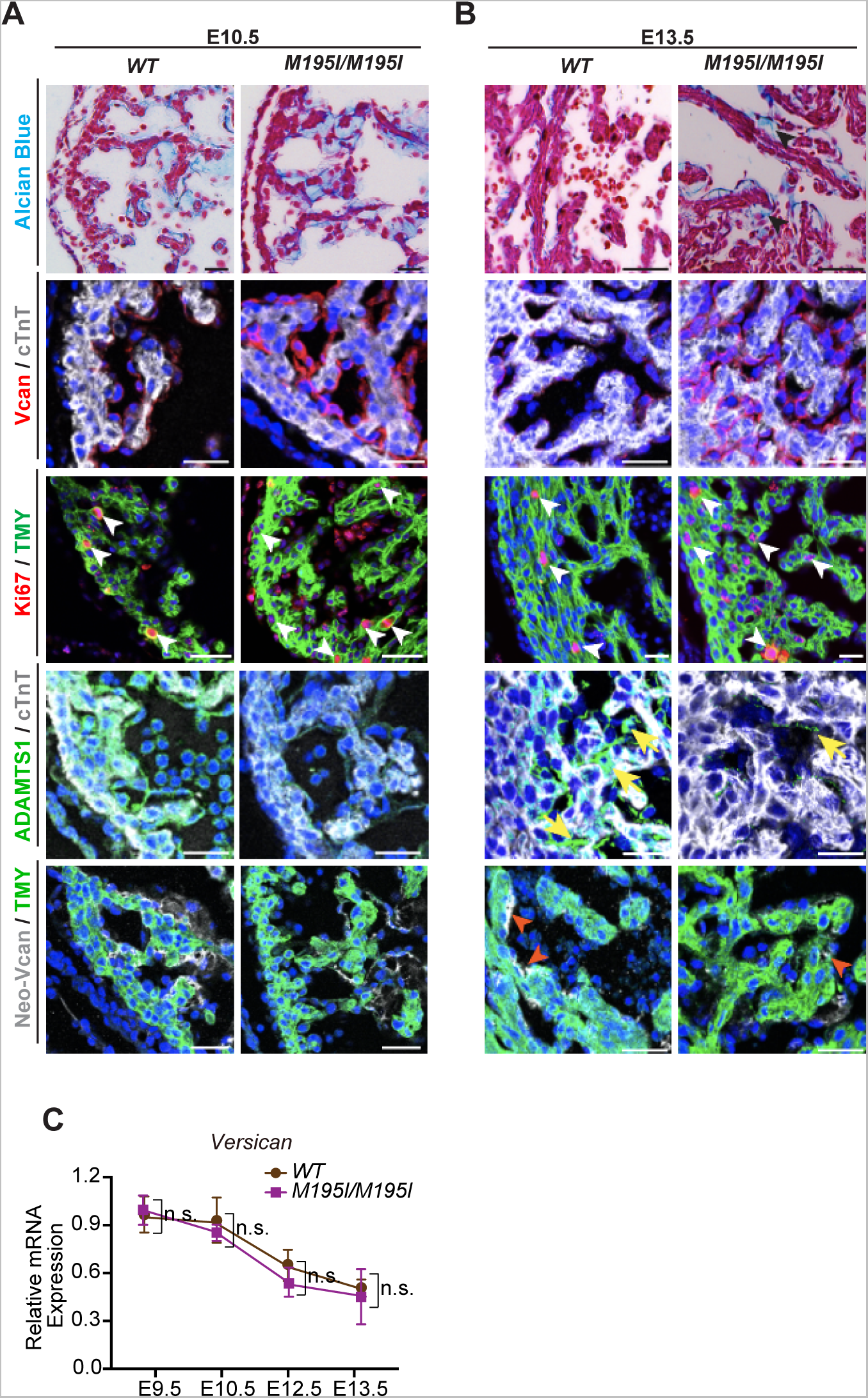
Cardiac ECM dynamics in *CHD4^M195I/M195I^* hearts are dysregulated. **A** and **B**, Representative images of Alcian Blue staining for extracellular matrix (ECM), immunofluorescent (Versican (Vcan), cardiac Troponin T (cTnT), and DAPI; or Ki67, Tropomyosin (TMY), and DAPI; or ADAMTS1, cTnT, and DAPI; or Neo-Versican (Neo-Vcan), TMY, and DAPI) stained paraffin sections of E10.5 (**A**) and E13.5 (**B**) hearts. Scale bars, 20 μm. **C**, RT-qPCR of *Versican* expression in WT and *CHD4^M195I/M195I^* hearts at indicated stages (n=3-5 nonpooled hearts per genotype per stage). *Pgk1* gene was used as the internal control, and *Versican* expression in E9.5 WT hearts was normalized as 1.0, expression of all other stages and *CHD4^M195I/M195I^* hearts were normalized to E9.5 WT hearts. Data are represented as mean ± SEM. Statistical significance was determined with one-way ANOVA (n.s., not significant (P>0.05)).

**Figure S11.**
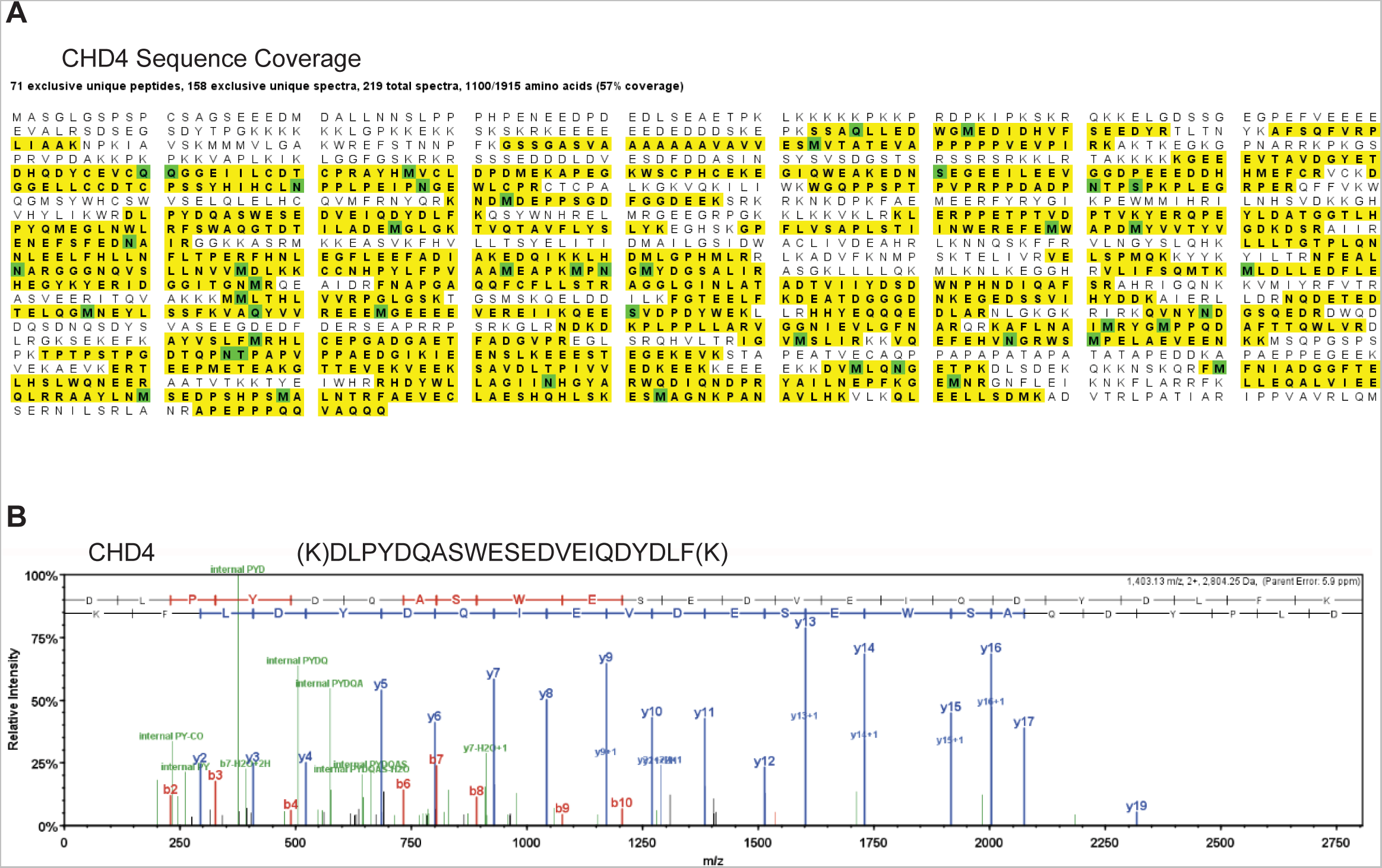
Validation of CHD4 immunoaffinity-MS in murine heart tissue. **A,** Sequence coverage of CHD4 from immune-isolates of E10.5 embryonic hearts (1100/1915 amino acids identified). **B,** Identification of CHD4 in immune-isolates of CHD4, as shown by CID MS/MS analysis of representative peptides.

**Figure S12.**
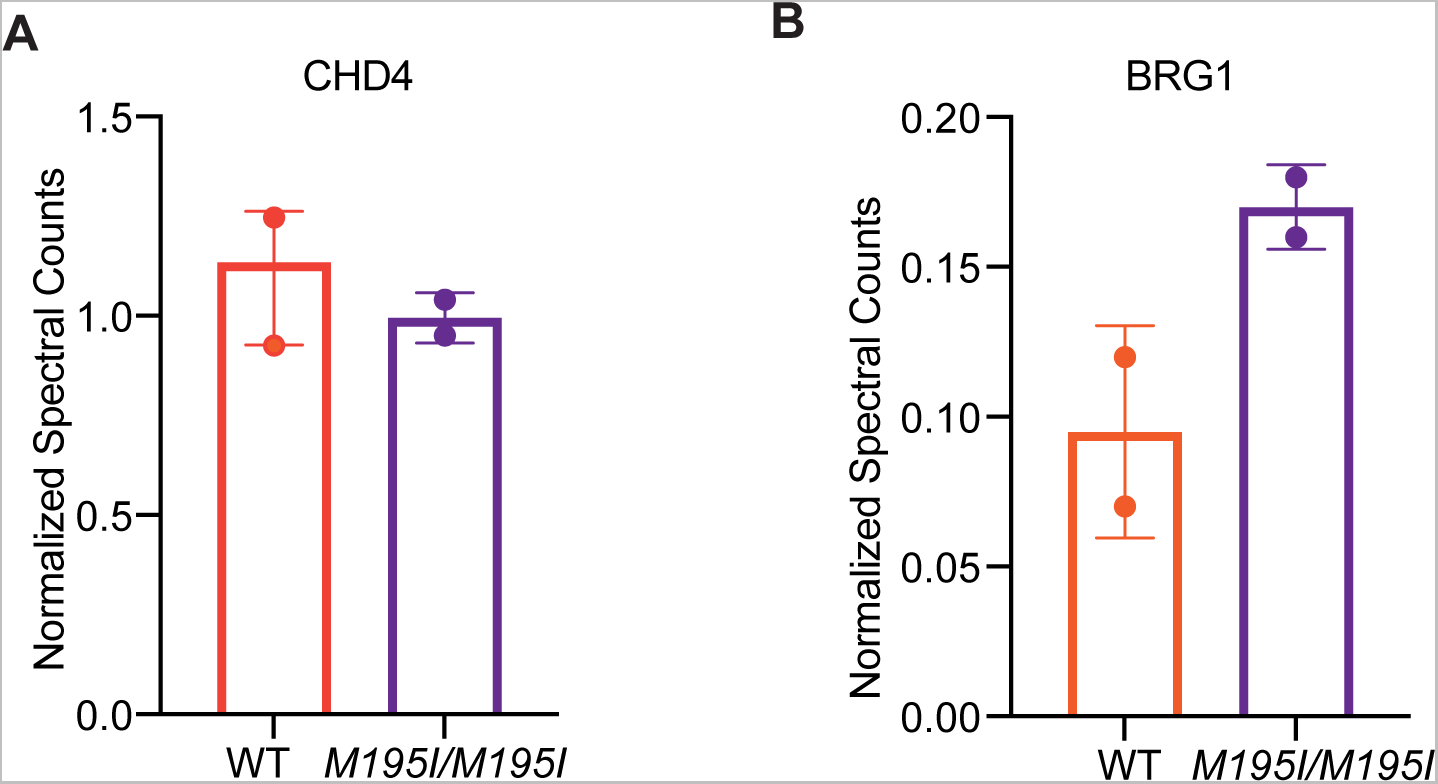
Normalized spectra count from CHD4-IP/MS at E13.5. **A** and **B**, CHD4 and BRG1 protein spectra counts normalized to CHD4 in CHD4-IP/MS results from WT and *CHD4^M195I/M195I^* hearts.

### Supplemental Methods

### Mice

The *CHD4^M195I^* mouse line was generated at the Animal Models Core of the University of North Carolina at Chapel Hill. Briefly, CAS9 guide RNAs flanking the coding sequences of the *Chd4* gene were identified using Benchling software. Four guide RNAs (Supplemental Table 1) at each end of the target sequence were selected for activity testing. This mutation introduced a Bcl1 restriction site for genotyping. RNAs were cloned into a T7 promoter vector followed by in vitro transcription and spin column purification. Functional testing was performed by transfecting a mouse embryonic fibroblast cell line with guide RNA and CAS9 protein. The guide RNA target site was amplified from transfected cells and analyzed by T7 Endonuclease 1 assay. C57BL/6J zygotes were electroporated with 1.2 μM CAS9 protein, 94.5 ng/μl each guide RNA and 400 ng/μl donor oligonucleotide and implanted in recipient pseudopregnant females. Resulting pups were screened by PCR followed by Bcl1 restriction digest for the presence of a mutant allele. Male founders with the correct deletion were mated to wild-type C57BL/6J females for germline transmission of the deletion allele. The founding *CHD4^M195I/+^* males were bred to wild-type mice for two generations, and the genotypes of *CHD4^M195I/+^* founding males and all F2 offspring were confirmed by sequencing and PCR/restriction digests. F2 mice (*CHD4^M195I/+^*) were intercrossed, and hearts derived from homozygous, heterozygous, and wild-type littermates were collected at desired developmental stages for specific assays, the breeding strategy is consistent across all assays unless where is specified.

Genotypes of the mice were confirmed by polymerase chain reaction (PCR) analysis using embryonic yolk sac or tail extracts. The amplicons were then purified with the QIAquick PCR Purification Kit (QIAGEN), and the purified amplicons were digested with the Bcl1 restriction enzyme at 50 °C for two hours. The purified amplicons were also sequenced and confirmed by sequencing. Primers for PCR and sequencing are listed in Supplemental Table 1.

All experiments were approved by the Institutional Animal Care and Use Committee at the University of North Carolina and conformed to the Guide for the Care and Use of Laboratory Animals.

### Histological Sectioning and Immunohistochemistry

The body torso containing hearts (for stages of E9.5 through E14.5), or the isolated hearts (for stages older than E14.5), or the *ex vivo* cultured hearts, were fixed in 4% PFA and paraffin-embedded. For the WGA staining, before fixation in 4% PFA, beating hearts were perfused with 10 mM KCl until the hearts were arrested. Paraffin sections (10 μm) were dewaxed, rehydrated, and stained with hematoxylin and eosin (H&E) using standard protocols^1^. Histology sections were imaged on an Olympus BX61 microscope. Immunohistochemistry staining and imaging were performed as previously reported^2^. Briefly, antigen retrieval was performed for the dewaxed. Rehydrated sections in boiled antigen retrieval buffer (10mM Sodium Citrate, 0.05% Tween 20, pH 6.0) for 20 min, cooled down sections were blocked with blocking buffer (10% NGS (normal goat serum; Sigma), 1% Triton X-100 in PBS) at R.T. for one hour, and then incubated with desired primary antibody solution (antibodies diluted in 10% blocking buffer supplemented with 5% donkey serum; detailed information for antibodies and dilution ratios were listed in Supplemental Table 2). overnight in a humidified chamber at 4°C. The next day, sections were washed three times with PBST (0.1% Tween 20 in PBS) and then incubated with secondary antibody solution (fluorescence-conjugated secondary antibodies diluted in PBST) for one hour at R.T., after washing with PBST three times, sections were counterstained with DAPI and mounted in Lab Vision PermaFluor Aqueous Mounting Medium (Fisher). Digital images were used for measurement using Fiji/ImageJ (NIH) software.

### Alcian Blue Staining

Alcian Blue staining was performed as previously described^3^. Briefly, the paraffin-embedded sections (10 μm) were dewaxed and rehydrated, followed by Alcian Blue solution staining (Sigma) at room temperature (R.T.) for 30 min. Then the slides were rinsed in running water for 5 min and counterstained with Nuclear Fast Red solution (Abcam) at R.T. for 10 min. After being stained, the slides were dehydrated through ethanol gradients and cleared by xylenes. Stained slides were images on Olympus BX61 microscope. The area of Alcian Blue-positive staining was measured using ImageJ (NIH) software.

### *In-situ* Proximity Ligation Assay

Proximity ligation assay (PLA) was performed as previously described with appropriate modifications^1^. Briefly, paraffin sections were dewaxed and rehydrated, followed by antigen retrieval treatment. Slides were blocked with 1% NGS (normal goat serum, Sigma)/1% Triton X-100/PBS at R.T. for 1 hour, rabbit anti-CHD4 (Abcam) and mouse anti-BRG1 (Santa Cruz) antibodies were probed at 4 C overnight. The next day, slides were washed with wash buffer and then were sequentially incubated with PLA probes for 1 hr at 37 °C, with ligase for 30 min at 37 °C and with polymerase for 100 min at 37 °C. After the final wash, mount the slides with a slide using PLA mounting medium with DAPI (PLA kit was purchased from Sigma). Images were captured on a Zeiss LSM 700 laser scanning confocal microscope.

### EdU Incorporation

Detection of proliferating cells by EdU incorporation assay was performed as previously described^4^. Briefly, time-mated females were subjected to intraperitoneal (i.p.) injection with 250 μg EdU 3 h before sacrifice. Embryos were recovered at E18.5, fixed with PFA and embedded for paraffin-sectioning. EdU staining was performed according to the manufacturer’s instructions. Antigen retrieval was performed, and co-stained antibodies were probed before applying the EdU detection cocktail. Eight or six coronal sections were analyzed for statistical analysis comprising two sections from four WT or three *CHD4^M195I/M195I^* embryos. The number of EdU^+^ cardiomyocytes among total cardiomyocytes (TMY^+^ cells) was counted from both ventricles.

### GW501516 Treatment of Mice

Plugged female mice were subjected to i.p. injection with GW501516 (Sigma; 3 mg/kg/day, dissolved in saline containing 0.5% DMSO) at E9.5, E10.5, and E11.5. Embryonic hearts were dissected on E12.5 and fixed in 4% PFA and paraffin-embedded. Three *M195I/+* dams were treated with GW501516, and four WT and *M195I/M195I* embryos were dissected and examined.

### *Ex vivo* Culture of Embryonic Heart Ventricles

*Ex* vivo culture of embryonic heart ventricles was performed as previously described with appropriate modifications^5, 6^. Briefly, plugged female mice were dissected at E11.5 and were cut through the middle line of the ventricles, the right ventricles were cultured in a small hole that was created on the low-melting point agarose gel; recombinant ADAMTS1 protein (Abcam; 5 μg/ml; BSA was used as control) was incubated with the Affi-Gel beads (Bio-Rad) at R.T. for two hours, and then the ADAMTS1 bead grafts were embedded beneath the inner side of the cultured ventricles for 24 hours. Explant ventricles were cultured in EdU-containing media (DMEM high glucose, 10% (vol/vol) FBS, 1% (vol/vol) MEM-NEAA (Thermo), 100 U/ml penicillin, 100 μg/ml streptomycin and 50 μM 2-mercaptoethanol) for the first 12 hours, and then cultured with EdU-free media for another 12 hours. Only beating ventricles were collected and fixed for following histological examinations, any non-beating explants should be discarded.

### Isolation of Cardiomyocytes from Embryonic Hearts

Isolation of cardiomyocytes from embryonic hearts was performed as previously described^7^ with appropriate modifications. Briefly, time-mated female mice were dissected at E18.5 and embryonic hearts were stored in individual tubes and were rinsed with pre-warmed PBS and culture media. 1 ml of trypsin-EDTA (0.05%) (Thermo Fisher) was added into each tube to digest hearts at 37 °C for 10 min. Discard the solution from this incubation, then add 1 ml of fresh trypsin-EDTA (0.05%) into each tube, break up the hearts with #19G needles, and then incubate at 37 °C for 10 min by rotating, after this incubation, collect the solution (excluding the tissues) to a new tube; repeat the trypsinization for six times, all solutions that contain dissociated cells were combined and spun at 1000 rpm for 10 min at R.T., the supernatant was aspirated and the cell pellet was resuspended with 3 ml of culture media. The resuspended cells were seeded in a 6-cm dish for 1 hour to allow those non-cardiomyocytes to attach to the plate while most cardiomyocytes were still in the culture media. Collect the culture media containing cardiomyocytes and count cell numbers with a hemocytometer for each sample.

### Flow Cytometry

EdU was injected into the pregnant dam at E18.5, 2 hours later, embryonic hearts were dissected and dissociated as described above. Dissociated cardiac cells were fixed with 4% PFA/PBS at room temperature for 15 min, and then quenched with 1% BSA/PBS. Discard the liquid and then block cells with 1% BSA/PBS at 4 °C overnight. The next day, spin cells down and then probe primary antibody (anti-TMY) at room temperature for 1 hour followed by three washes and secondary antibody probing at room temperature for 1 hour. EdU detection, flow cytometry gating strategy, and analysis were performed as previously described^8^.

### Immuno-purification

Frozen hearts from E18.5 (9 hearts were pooled as one replicate, two replicates for each genotype) or E12.5 (20 hearts were pooled as one replicate, two replicates for each genotype) embryos were homogenized using a mortar and pestle in liquid nitrogen. Nuclei were prepared as previously described^9^. Isolated nuclei were resuspended in optimized lysis buffer (20 mM K-HEPES pH 7.4, 0.11 M KOAC, 2 mM MgCl_2_, 0.1% Tween 20, 1 μM ZnCl_2_, 1 mM CaCl_2_, 0.5% Triton-X 100, 150 mM NaCl, protease inhibitor (Sigma), phosphatase inhibitor (Sigma)). Samples were homogenized using a Polytron (Kinematica) followed by universal nuclease treatment (Pierce). The cleared lysates were incubated with the anti-CHD4 antibody at 4 °C overnight, while the Protein A/G dynabeads (Thermo) were blocked with 75 mg/ml BSA (Sigma) at 4 °C overnight. The next day, discard the BSA solution from the beads, and incubate the blocked beads with the lysate-antibody mixture at 4 °C for two hours. After 2-hour incubation, discard the supernatant and wash the beads with wash buffer (same components as the lysis buffer without protease/phosphatase inhibitors) six times; proteins were eluted at 95 °C for 10 min. The immune-isolated proteins were subjected to Western Blot or Mass Spectrometry analysis.

### Western Blot

Western blots were blocked with 5% milk/TBST at room temperature for one hour, and then probed with following primary antibodies overnight at 4°C: rabbit anti-CHD4 (Active Motif), mouse anti-GAPDH (Millipore), mouse anti-BRG1 (Santa Cruz). After being rinsed, blots were probed with the HRP-conjugated secondary antibodies (Jackson Immunoresearch) for one hour at room temperature. Antibody-antigen complexes were visualized using an ECL Western Blotting Analysis System (Amersham).

### Proteomic Analysis of CHD4 Affinity Purifications

Mass spectrometry was conducted as previously reported^1^. Briefly, immunoisolated proteins were resolved (∼ 4 cm) by SDS-PAGE and visualized by Coomassie blue. Each lane was subjected to in-gel digestion with trypsin and analyzed by nano liquid chromatography coupled to tandem mass spectrometry as previously reported (Robbe et al., 2022). Tandem mass spectra were extracted by Proteome Discoverer (ThermoFisher Scientific, ver 1.4), and searched with the SEQUEST algorithm against a theoretical tryptic peptide database generated from the forward or reverse entries of the mouse UniProt-SwissProt protein sequence database (2013/08) and common contaminants (total of 43, 007 sequences). SEQUEST search results were analyzed by Scaffold (version 4.6.1, Proteome Software Inc) using the LFDR scoring scheme to calculate peptide and protein probabilities. Peptide and protein probabilities thresholds were selected to achieve ≤ 1% FDR at the peptide level based on LFDR modeling and at the protein level, based on the number of proteins identified as hits to the reverse database. The spectral counts assigned to proteins that satisfied these criteria and had a minimum of two unique peptides were exported to Excel for data processing.

### Transmission Electron Microscopy

The TEM examination was performed as previously described^2^. Briefly, E18.5 embryos were collected and fixed in 2% formaldehyde/2.5% glutaraldehyde in 0.15M sodium phosphate buffer, pH 7.4, overnight at 4°C. Following several washes in sodium phosphate buffer, samples were post-fixed for 1 hour in 1% buffered osmium tetroxide, dehydrated through a graded series of ethanol and embedded in PolyBed 812 epoxy resin (Polysciences). The cut sections were observed using an LEO EM-910 transmission electron microscope (LEO Electron Microscopy Inc.), accelerating voltage of 80kV and micrographs were taken using a Gatan Orius SC1000 CCD Camera.

### Echocardiography

Echocardiography on adult mice was performed as previously described^10^. Noninvasive M-mode echocardiography was performed on E18.5 embryos as previously described^11–13^. Briefly, pregnant female mice were anesthetized with 2% isoflurane until they became nonresponsive. Fix and monitor the pregnant mice on the measurement platform appropriately, and then the hair removal cream was applied to the abdomen from chest to bladder to remove the hair quickly but gently. Press down on the naked abdomen gently to locate the embryos. Slowly and lightly spread them out to have most of embryos in a single layer under the abdominal surface. Mark each embryo on the naked abdomen of the dam with a permanent marker with their anterior/posterior and dorsal/ventral directions, take a picture for these marks for matching genotypes and acquired data. Place a small amount of prewarmed ultrasound gel on the naked abdomen and spread it evenly. Hold the probe in contact with the thick gel layer and gradually move the probe toward the skin while looking for the beating heart. Once the beating heart is visualized on the screen, begin to acquire videos. After examining all marked embryos, sacrifice the female mice by applying CO_2_ and cervical dislocation, incise the skin and muscle layer of the abdomen longitudinally carefully, take every embryo out from the uterine horn and label them with marked numbers by referring to the picture that was taken earlier. Dissect and store the embryonic hearts for other assays (i.e., IP/MS, IHC), and reserve a small tissue for genotyping. Analyze the data and match the data to genotypes.

### RNA Extraction, RNA-Sequencing, and RT-qPCR

RNA from individual embryonic hearts was extracted using the RNAqueous-Micro total RNA isolation kit following the manufacturer’s recommended protocols (Invitrogen), as previously described^1^. The concentration and quality of purified RNA were assessed by TapeStation (Agilent). RNA integrity numbers (RIN) ranged from 8.0 to 10.0. For RNA extracted from E18.5 hearts, purified poly-A RNA that had undergone two rounds of oligo-dT selection was converted into cDNA and used to generate RNA-seq libraries. Libraries were sequenced (75-bp paired-end reads; Illumina HiSeq 2500) to a target depth of >30 million reads. Reads were aligned to the mm10 reference genome using STAR via the bcbio-nextgen RNA sequencing pipeline. RNAseq analysis was performed using DESeq2 (DESeq2_1.18.1) in R (3.6.2). Genes with a >0.5 log2 (fold change) and an adjusted P-value < 0.05 were considered statistically significant. Pathway enrichment analysis was performed with the Gene Set Enrichment Analysis (GSEA) tool, a computational method that detects modest but coordinated changes in the expression of groups of functionally related genes^14, 15^.

For RT-qPCR, cDNA synthesis was performed using random hexamers and SuperScript IV reverse transcriptase (Invitrogen). Quantitative PCR was performed using PowerUP SYBR Green master mix (Thermo) at standard cycling conditions on a QuantStudio 7 Flex instrument (Thermo) with the primers in Supplemental Table 1.

### Chromatin Immunoprecipitation (ChIP)-sequencing, Analysis, and ChIP-qPCR

Hearts from WT, *CHD4^M195I/+^*, and *CHD4^M195I/M195I^* embryos at E12.5 (15 hearts per biological replicate, two biological replicates per genotype) were fixed for chromatin immunoprecipitation as described previously^2^. Briefly, hearts were dissected in cold PBS and crosslinked with 1% PFA/PBS for 10 minutes at room temperature. Fixation was quenched with 125 mM glycine. Hearts were washed twice with cold PBS, snap frozen and stored at −80°C. Hearts were prepared for ChIP as described^2^ with some modifications. Frozen hearts were resuspended and washed in Hypotonic buffer (10 mM HEPES-NaOH pH 7.9, 10 mM KCl, 1.5mM MgCl_2_, 340 mM sucrose, 10% glycerol, 0.1% Triton-X 100, protease inhibitors (Sigma)) and then resuspended and dounced 20 times on ice in shearing buffer (10 mM Tris-HCl pH 8.0, 1 mM EDTA, 0.5 mM EGTA, 0.5 mM PMSF, 5 mM NaBut, 0.1% SDS, protease inhibitors). Homogenates were transferred and sonicated in a Covaris S220 with the following settings: power peak 175, duty factor 10, 200 counts for 1200 seconds at 4°C. Following sonication, samples were centrifuged at 20000 g at 4°C for 10 min. The soluble chromatin fraction was diluted 1:1 with 2×IP buffer (20 mM Tris-HCl pH 8.0, 300 mM NaCl, 2 mM EDTA, 20% glycerol, 1% Triton-X 100, 5 mM NaBut, 0.5 mM PMSF, protease inhibitors). 10% of the input was stored at −80°C for input library sequencing. 700 μl of the chromatin fraction was mixed with 3 μg rabbit anti-CHD4 antibody (Abcam) and rotated overnight at 4°C. 20 μl of each Protein A (Thermo) and Protein G Dynabeads (Thermo) were blocked overnight at 4°C with 75 mg/ml BSA (final concentration: 5 mg/ml; Sigma). The next day, blocked beads were rotated with the chromatin, immunoprecipitated for 3 hrs at 4°C, then washed orderly with low salt buffer (20 mM Tris-HCl pH 8.0, 150 mM NaCl, 2 mM EDTA, 1% Triton-X 100, 0.1% SDS), high salt buffer (20 mM Tris-HCl pH 8.0, 300 mM NaCl, 2 mM EDTA, 1% Triton-X 100, 0.1% SDS), LiCl buffer (20 mM Tris-HCl pH 8.0, 250 mM LiCl, 2 mM EDTA, 1% NP-40, 1%/vol NaDeoxycholate), TE10/1 buffer (10 mM Tris-HCl pH 8.0, 1 mM EDTA). Chromatin was eluted with freshly prepared elution buffer (1% SDS, 0.1 M NaHCO_3_, 5 mM DTT) for 1 hr at 65°C. Eluted chromatin was incubated with 6 µl of 5 M NaCl and 0.5 µl of RNase cocktail for 4 hrs at 65°C to reverse crosslink, then 2 µl of 1 M Tris-HCl pH 6.8 and 2 µl of 20 mg/ml Proteinase K were added and incubated for 1 hr at 65°C. DNA was purified with Ampure XP beads (Beckman coulter). Libraries were prepared using the ThruPLEX-FDPrep Kit. Libraries were sequenced using the HiSeq4000 at the UNC High-throughput Sequencing Facility (HTSF) with 50 base pair paired-end reads per sample.

Purified input and CHD4 ChIP DNA used for NGS library preparation was also used for confirmation by qPCR. Briefly, 10% input and CHD4 ChIP DNA were diluted 1:10 for each reaction. Quantitative PCR was performed using PowerUP SYBR Green Master Mix (Thermo) at standard cycling conditions and reaction composition on a QuantStudio 7 Flex instrument (Thermo) with primers listed in Supplemental Table 1.

### Quantification and Statistical Analysis

Ventricular and IVS thickness were measured with Fiji/ImageJ software. Compact myocardium thickness was measured from the base of trabeculae to epicardium, and trabecular myocardium thickness was determined by Endomucin staining. For an individual heart, all measurements were averaged at values of 20-30 different positions on its sections.

For cardiomyocyte proliferation, cardiomyocyte number was counted manually (overlay of TMY (or cTnT) and DAPI). EdU^+^/TMY^+^ (or EdU^+^/cTnT^+^, or Ki67^+^/TMY^+^) nuclei (proliferative cardiomyocytes) were identified by merging EdU or Ki67 images with TMY or cTnT images, and using Fiji/ImageJ color threshold function to select for the merged color signal generated by overlaying TMY or cTnT signal and EdU or Ki67 signal. Double positive stained cells were counted with Fiji/ImageJ cell count function.

All data were presented as mean ± SEM and statistically analyzed using the Prism 9 software. Numbers of independent biological replicates were provided in the captions of the figures. The two-tailed unpaired Student’s t-test was used to compare the mean difference between 2 groups. One-way or two-way ANOVA was used to compare the mean from ≥3 groups. If ANOVA analysis showed a significant difference, then the Tukey multiple comparison tests was applied for post hoc analysis to detect the pairwise difference while adjusting for multiplicity. P<0.05 was considered statistically significant.

## Supplemental Tables

**Supplemental Table 1:**
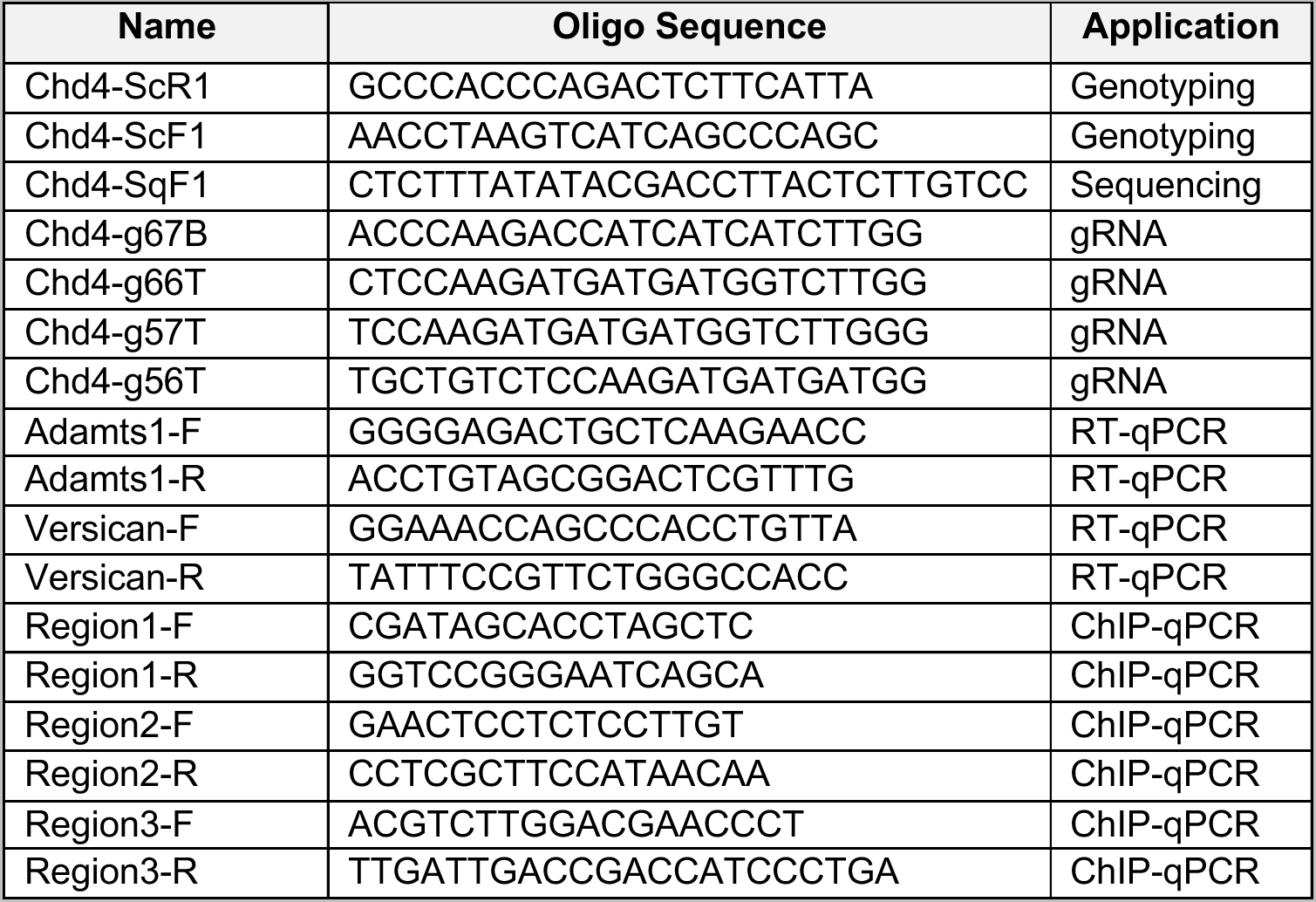
Oligo sequences used in this paper.

**Supplemental Table 2:**
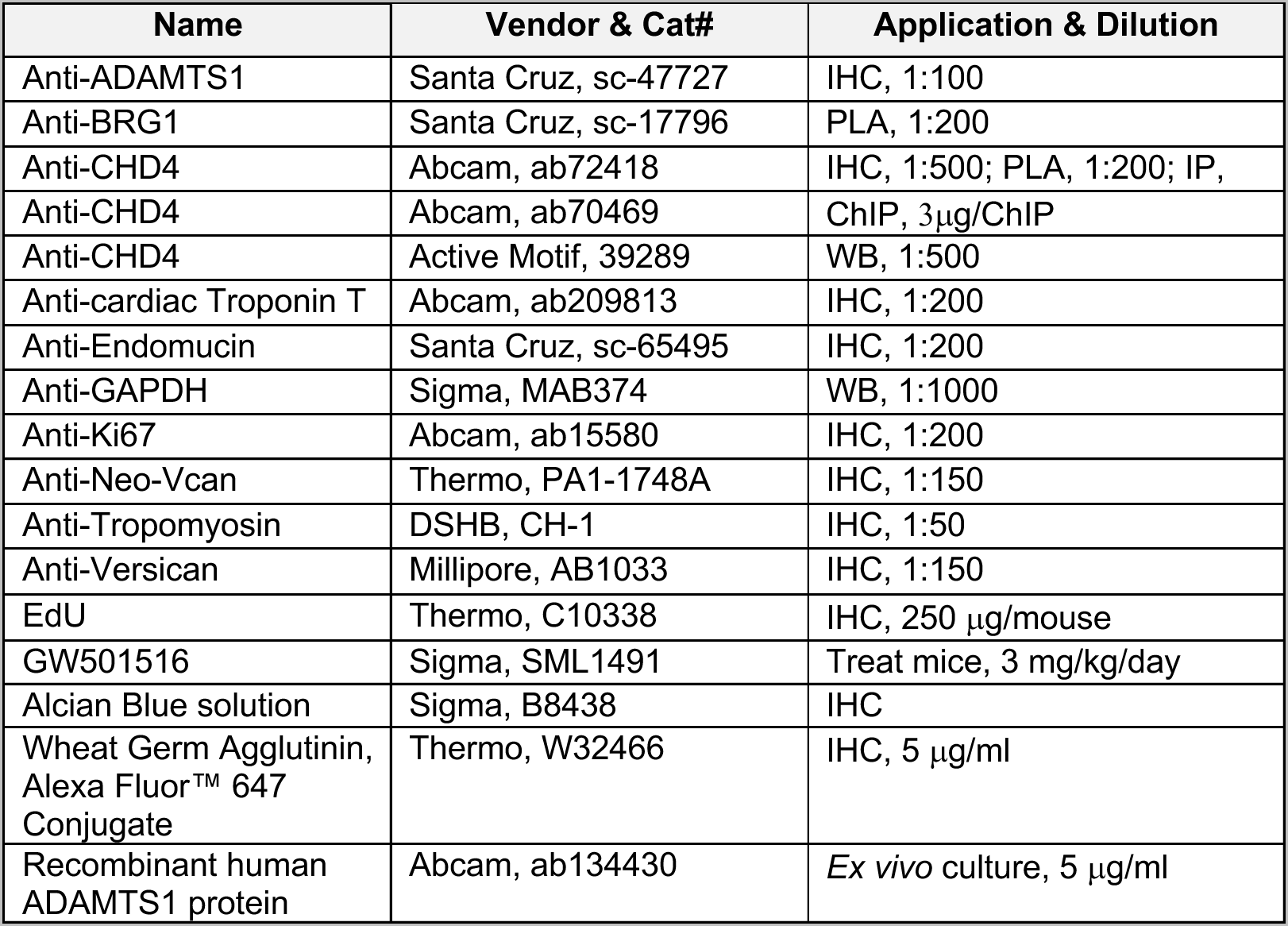
Critical reagents used in this paper.

| Name | Vendor & Cat# | Application & Dilution |
| --- | --- | --- |
| Anti-ADAMTS1 | Santa Cruz, sc-47727 | IHC, 1:100 |
| Anti-BRG1 | Santa Cruz, sc-17796 | PLA, 1:200 |
| Anti-CHD4 | Abcam, ab72418 | IHC, 1:500; PLA, 1:200; IP, |
| Anti-CHD4 | Abcam, ab70469 | ChIP, 3µg/ChIP |
| Anti-CHD4 | Active Motif, 39289 | WB, 1:500 |
| Anti-cardiac Troponin T | Abcam, ab209813 | IHC, 1:200 |
| Anti-Endomucin | Santa Cruz, sc-65495 | IHC, 1:200 |
| Anti-GAPDH | Sigma, MAB374 | WB, 1:1000 |
| Anti-Ki67 | Abcam, ab15580 | IHC, 1:200 |
| Anti-Neo-Vcan | Thermo, PA1-1748A | IHC, 1:150 |
| Anti-Tropomyosin | DSHB, CH-1 | IHC, 1:50 |
| Anti-Versican | Millipore, AB1033 | IHC, 1:150 |
| EdU | Thermo, C10338 | IHC, 250 µg/mouse |
| GW501516 | Sigma, SML1491 | Treat mice, 3 mg/kg/day |
| Alcian Blue solution | Sigma, B8438 | IHC |
| Wheat Germ Agglutinin, Alexa Fluor™ 647 Conjugate | Thermo, W32466 | IHC, 5 µg/ml |
| Recombinant human ADAMTS1 protein | Abcam, ab134430 | Ex vivo culture, 5 µg/ml |

## Supplemental Videos

Supplemental Videos S1-S3: In utero echocardiography of E18.5 WT, *CHD4^M195I/+^*, and *CHD4^M195I/M195I^* embryos, related to Figure 3.

Supplemental Videos S4 and S5: Echocardiography of adult WT and *CHD4^M195I/+^* mice, related to Figure S5.

## References

1. Finsterer J, Stollberger C and Towbin JA. Left ventricular noncompaction cardiomyopathy: cardiac, neuromuscular, and genetic factors. Nat Rev Cardiol. 2017;14:224–237.

2. Ramakumar V, Sharma V and Mishra S. Left ventricular non-compaction. Eur Heart J. 2021;42:2398.

3. Fazio G, Lunetta M, Grassedonio E, Gullotti A, Ferro G, Bacarella D, Lo Re G, Novo G, Massimo M, Maresi E and Novo S. Noncompaction of the right ventricle. Pediatr Cardiol. 2010;31:576–8.

4. Ranganathan A, Ganesan G, Sangareddi V, Pillai AP and Ramasamy A. Isolated noncompaction of right ventricle--a case report. Echocardiography. 2012;29:E169–72.

5. Syed MP, Doshi A, Pandey D, Kate Y and Harizi R. A rare case of biventricular non-compaction. BMJ Case Rep. 2020;13.

6. Weiford BC, Subbarao VD and Mulhern KM. Noncompaction of the ventricular myocardium. Circulation. 2004;109:2965–71.

7. Klaassen S, Probst S, Oechslin E, Gerull B, Krings G, Schuler P, Greutmann M, Hurlimann D, Yegitbasi M, Pons L, Gramlich M, Drenckhahn JD, Heuser A, Berger F, Jenni R and Thierfelder L. Mutations in sarcomere protein genes in left ventricular noncompaction. Circulation. 2008;117:2893–901.

8. Towbin JA. Ion channel dysfunction associated with arrhythmia, ventricular noncompaction, and mitral valve prolapse: a new overlapping phenotype. J Am Coll Cardiol. 2014;64:768–71.

9. Ashraf H, Pradhan L, Chang EI, Terada R, Ryan NJ, Briggs LE, Chowdhury R, Zarate MA, Sugi Y, Nam HJ, Benson DW, Anderson RH and Kasahara H. A mouse model of human congenital heart disease: high incidence of diverse cardiac anomalies and ventricular noncompaction produced by heterozygous Nkx2-5 homeodomain missense mutation. Circ Cardiovasc Genet. 2014;7:423–433.

10. Gan P, Wang Z, Morales MG, Zhang Y, Bassel-Duby R, Liu N and Olson EN. RBPMS is an RNA-binding protein that mediates cardiomyocyte binucleation and cardiovascular development. Dev Cell. 2022;57:959–973 e7.

11. Pantazis AA and Elliott PM. Left ventricular noncompaction. Curr Opin Cardiol. 2009;24:209–13.

12. Sedmera D, Pexieder T, Vuillemin M, Thompson RP and Anderson RH. Developmental patterning of the myocardium. Anat Rec. 2000;258:319–37.

13. Del Monte-Nieto G, Ramialison M, Adam AAS, Wu B, Aharonov A, D’Uva G, Bourke LM, Pitulescu ME, Chen H, de la Pompa JL, Shou W, Adams RH, Harten SK, Tzahor E, Zhou B and Harvey RP. Control of cardiac jelly dynamics by NOTCH1 and NRG1 defines the building plan for trabeculation. Nature. 2018;557:439–445.

14. D’Amato G, Luxan G, del Monte-Nieto G, Martinez-Poveda B, Torroja C, Walter W, Bochter MS, Benedito R, Cole S, Martinez F, Hadjantonakis AK, Uemura A, Jimenez-Borreguero LJ and de la Pompa JL. Sequential Notch activation regulates ventricular chamber development. Nat Cell Biol. 2016;18:7–20.

15. Luxan G, Casanova JC, Martinez-Poveda B, Prados B, D’Amato G, MacGrogan D, Gonzalez-Rajal A, Dobarro D, Torroja C, Martinez F, Izquierdo-Garcia JL, Fernandez-Friera L, Sabater-Molina M, Kong YY, Pizarro G, Ibanez B, Medrano C, Garcia-Pavia P, Gimeno JR, Monserrat L, Jimenez-Borreguero LJ and de la Pompa JL. Mutations in the NOTCH pathway regulator MIB1 cause left ventricular noncompaction cardiomyopathy. Nat Med. 2013;19:193–201.

16. Wade PA, Gegonne A, Jones PL, Ballestar E, Aubry F and Wolffe AP. Mi-2 complex couples DNA methylation to chromatin remodelling and histone deacetylation. Nat Genet. 1999;23:62–6.

17. Xue Y, Wong J, Moreno GT, Young MK, Cote J and Wang W. NURD, a novel complex with both ATP-dependent chromatin-remodeling and histone deacetylase activities. Mol Cell. 1998;2:851–61.

18. Zhang Y, LeRoy G, Seelig HP, Lane WS and Reinberg D. The dermatomyositis-specific autoantigen Mi2 is a component of a complex containing histone deacetylase and nucleosome remodeling activities. Cell. 1998;95:279–89.

19. O’Shaughnessy-Kirwan A, Signolet J, Costello I, Gharbi S and Hendrich B. Constraint of gene expression by the chromatin remodelling protein CHD4 facilitates lineage specification. Development. 2015;142:2586–97.

20. Hung H, Kohnken R and Svaren J. The nucleosome remodeling and deacetylase chromatin remodeling (NuRD) complex is required for peripheral nerve myelination. J Neurosci. 2012;32:1517–27.

21. Kashiwagi M, Morgan BA and Georgopoulos K. The chromatin remodeler Mi-2beta is required for establishment of the basal epidermis and normal differentiation of its progeny. Development. 2007;134:1571–82.

22. Williams CJ, Naito T, Arco PG, Seavitt JR, Cashman SM, De Souza B, Qi X, Keables P, Von Andrian UH and Georgopoulos K. The chromatin remodeler Mi-2beta is required for CD4 expression and T cell development. Immunity. 2004;20:719–33.

23. Yoshida T, Hazan I, Zhang J, Ng SY, Naito T, Snippert HJ, Heller EJ, Qi X, Lawton LN, Williams CJ and Georgopoulos K. The role of the chromatin remodeler Mi-2beta in hematopoietic stem cell self-renewal and multilineage differentiation. Genes Dev. 2008;22:1174–89.

24. Yoshida T, Hu Y, Zhang Z, Emmanuel AO, Galani K, Muhire B, Snippert HJ, Williams CJ, Tolstorukov MY, Gounari F and Georgopoulos K. Chromatin restriction by the nucleosome remodeler Mi-2beta and functional interplay with lineage-specific transcription regulators control B-cell differentiation. Genes Dev. 2019;33:763–781.

25. Homsy J, Zaidi S, Shen Y, Ware JS, Samocha KE, Karczewski KJ, DePalma SR, McKean D, Wakimoto H, Gorham J, Jin SC, Deanfield J, Giardini A, Porter GA, Jr., Kim R, Bilguvar K, Lopez-Giraldez F, Tikhonova I, Mane S, Romano-Adesman A, Qi H, Vardarajan B, Ma L, Daly M, Roberts AE, Russell MW, Mital S, Newburger JW, Gaynor JW, Breitbart RE, Iossifov I, Ronemus M, Sanders SJ, Kaltman JR, Seidman JG, Brueckner M, Gelb BD, Goldmuntz E, Lifton RP, Seidman CE and Chung WK. De novo mutations in congenital heart disease with neurodevelopmental and other congenital anomalies. Science. 2015;350:1262–6.

26. Wilczewski CM, Hepperla AJ, Shimbo T, Wasson L, Robbe ZL, Davis IJ, Wade PA and Conlon FL. CHD4 and the NuRD complex directly control cardiac sarcomere formation. Proc Natl Acad Sci U S A. 2018;115:6727–6732.

27. Malecova B, Dall’Agnese A and Puri PL. Muscles cannot break a NuRDy heart. EMBO J. 2016;35:1600–2.

28. Gomez-Del Arco P, Perdiguero E, Yunes-Leites PS, Acin-Perez R, Zeini M, Garcia-Gomez A, Sreenivasan K, Jimenez-Alcazar M, Segales J, Lopez-Maderuelo D, Ornes B, Jimenez-Borreguero LJ, D’Amato G, Enshell-Seijffers D, Morgan B, Georgopoulos K, Islam AB, Braun T, de la Pompa JL, Kim J, Enriquez JA, Ballestar E, Munoz-Canoves P and Redondo JM. The Chromatin Remodeling Complex Chd4/NuRD Controls Striated Muscle Identity and Metabolic Homeostasis. Cell Metab. 2016;23:881–92.

29. Robbe ZL, Shi W, Wasson LK, Scialdone AP, Wilczewski CM, Sheng X, Hepperla AJ, Akerberg BN, Pu WT, Cristea IM, Davis IJ and Conlon FL. CHD4 is recruited by GATA4 and NKX2-5 to repress noncardiac gene programs in the developing heart. Genes Dev. 2022;36:468–482.

30. Jin SC, Homsy J, Zaidi S, Lu Q, Morton S, DePalma SR, Zeng X, Qi H, Chang W, Sierant MC, Hung WC, Haider S, Zhang J, Knight J, Bjornson RD, Castaldi C, Tikhonoa IR, Bilguvar K, Mane SM, Sanders SJ, Mital S, Russell MW, Gaynor JW, Deanfield J, Giardini A, Porter GA, Jr., Srivastava D, Lo CW, Shen Y, Watkins WS, Yandell M, Yost HJ, Tristani-Firouzi M, Newburger JW, Roberts AE, Kim R, Zhao H, Kaltman JR, Goldmuntz E, Chung WK, Seidman JG, Gelb BD, Seidman CE, Lifton RP and Brueckner M. Contribution of rare inherited and de novo variants in 2,871 congenital heart disease probands. Nat Genet. 2017;49:1593–1601.

31. Pediatric Cardiac Genomics C, Gelb B, Brueckner M, Chung W, Goldmuntz E, Kaltman J, Kaski JP, Kim R, Kline J, Mercer-Rosa L, Porter G, Roberts A, Rosenberg E, Seiden H, Seidman C, Sleeper L, Tennstedt S, Kaltman J, Schramm C, Burns K, Pearson G and Rosenberg E. The Congenital Heart Disease Genetic Network Study: rationale, design, and early results. Circ Res. 2013;112:698–706.

32. Weiss K, Lazar HP, Kurolap A, Martinez AF, Paperna T, Cohen L, Smeland MF, Whalen S, Heide S, Keren B, Terhal P, Irving M, Takaku M, Roberts JD, Petrovich RM, Schrier Vergano SA, Kenney A, Hove H, DeChene E, Quinonez SC, Colin E, Ziegler A, Rumple M, Jain M, Monteil D, Roeder ER, Nugent K, van Haeringen A, Gambello M, Santani A, Medne L, Krock B, Skraban CM, Zackai EH, Dubbs HA, Smol T, Ghoumid J, Parker MJ, Wright M, Turnpenny P, Clayton-Smith J, Metcalfe K, Kurumizaka H, Gelb BD, Baris Feldman H, Campeau PM, Muenke M, Wade PA and Lachlan K. The CHD4-related syndrome: a comprehensive investigation of the clinical spectrum, genotype-phenotype correlations, and molecular basis. Genet Med. 2020;22:389–397.

33. Silva AP, Ryan DP, Galanty Y, Low JK, Vandevenne M, Jackson SP and Mackay JP. The N-terminal Region of Chromodomain Helicase DNA-binding Protein 4 (CHD4) Is Essential for Activity and Contains a High Mobility Group (HMG) Box-like-domain That Can Bind Poly(ADP-ribose). J Biol Chem. 2016;291:924–38.

34. Pan MR, Hsieh HJ, Dai H, Hung WC, Li K, Peng G and Lin SY. Chromodomain helicase DNA-binding protein 4 (CHD4) regulates homologous recombination DNA repair, and its deficiency sensitizes cells to poly(ADP-ribose) polymerase (PARP) inhibitor treatment. J Biol Chem. 2012;287:6764–72.

35. Stacey RB, Caine AJ, Jr. and Hundley WG. Evaluation and management of left ventricular noncompaction cardiomyopathy. Curr Heart Fail Rep. 2015;12:61–7.

36. Kawel N, Nacif M, Arai AE, Gomes AS, Hundley WG, Johnson WC, Prince MR, Stacey RB, Lima JA and Bluemke DA. Trabeculated (noncompacted) and compact myocardium in adults: the multi-ethnic study of atherosclerosis. Circ Cardiovasc Imaging. 2012;5:357–66.

37. Kim W, Seidah NG and Prat A. In utero measurement of heart rate in mouse by noninvasive M-mode echocardiography. J Vis Exp. 2013:e50994.

38. Touma M. Fetal Mouse Cardiovascular Imaging Using a High-frequency Ultrasound (30/45MHZ) System. J Vis Exp. 2018.

39. Mei L, Kedziora KM, Song EA, Purvis JE and Cook JG. The consequences of differential origin licensing dynamics in distinct chromatin environments. Nucleic Acids Res. 2022.

40. Guo Y and Pu WT. Cardiomyocyte Maturation: New Phase in Development. Circ Res. 2020;126:1086–1106.

41. Puente BN, Kimura W, Muralidhar SA, Moon J, Amatruda JF, Phelps KL, Grinsfelder D, Rothermel BA, Chen R, Garcia JA, Santos CX, Thet S, Mori E, Kinter MT, Rindler PM, Zacchigna S, Mukherjee S, Chen DJ, Mahmoud AI, Giacca M, Rabinovitch PS, Aroumougame A, Shah AM, Szweda LI and Sadek HA. The oxygen-rich postnatal environment induces cardiomyocyte cell-cycle arrest through DNA damage response. Cell. 2014;157:565–79.

42. Kannan S and Kwon C. Regulation of cardiomyocyte maturation during critical perinatal window. J Physiol. 2020;598:2941–2956.

43. Franco D, Lamers WH and Moorman AF. Patterns of expression in the developing myocardium: towards a morphologically integrated transcriptional model. Cardiovasc Res. 1998;38:25–53.

44. Lyons GE, Schiaffino S, Sassoon D, Barton P and Buckingham M. Developmental regulation of myosin gene expression in mouse cardiac muscle. J Cell Biol. 1990;111:2427–36.

45. Bersell K, Arab S, Haring B and Kuhn B. Neuregulin1/ErbB4 signaling induces cardiomyocyte proliferation and repair of heart injury. Cell. 2009;138:257–70.

46. Ahuja P, Perriard E, Perriard JC and Ehler E. Sequential myofibrillar breakdown accompanies mitotic division of mammalian cardiomyocytes. J Cell Sci. 2004;117:3295–306.

47. Neubauer S. The failing heart--an engine out of fuel. N Engl J Med. 2007;356:1140–51.

48. Lopaschuk GD and Jaswal JS. Energy metabolic phenotype of the cardiomyocyte during development, differentiation, and postnatal maturation. J Cardiovasc Pharmacol. 2010;56:130–40.

49. Calmettes G, John SA, Weiss JN and Ribalet B. Hexokinase-mitochondrial interactions regulate glucose metabolism differentially in adult and neonatal cardiac myocytes. J Gen Physiol. 2013;142:425–36.

50. DeLaughter DM, Bick AG, Wakimoto H, McKean D, Gorham JM, Kathiriya IS, Hinson JT, Homsy J, Gray J, Pu W, Bruneau BG, Seidman JG and Seidman CE. Single-Cell Resolution of Temporal Gene Expression during Heart Development. Dev Cell. 2016;39:480–490.

51. Lockhart M, Wirrig E, Phelps A and Wessels A. Extracellular matrix and heart development. Birth Defects Res A Clin Mol Teratol. 2011;91:535–50.

52. Camenisch TD, Spicer AP, Brehm-Gibson T, Biesterfeldt J, Augustine ML, Calabro A, Jr., Kubalak S, Klewer SE and McDonald JA. Disruption of hyaluronan synthase-2 abrogates normal cardiac morphogenesis and hyaluronan-mediated transformation of epithelium to mesenchyme. J Clin Invest. 2000;106:349–60.

53. Sandireddy R, Cibi DM, Gupta P, Singh A, Tee N, Uemura A, Epstein JA and Singh MK. Semaphorin 3E/PlexinD1 signaling is required for cardiac ventricular compaction. JCI Insight. 2019;4.

54. Stankunas K, Hang CT, Tsun ZY, Chen H, Lee NV, Wu JI, Shang C, Bayle JH, Shou W, Iruela-Arispe ML and Chang CP. Endocardial Brg1 represses ADAMTS1 to maintain the microenvironment for myocardial morphogenesis. Dev Cell. 2008;14:298–311.

55. Yamamura H, Zhang M, Markwald RR and Mjaatvedt CH. A heart segmental defect in the anterior-posterior axis of a transgenic mutant mouse. Dev Biol. 1997;186:58–72.

56. Chan CK, Rolle MW, Potter-Perigo S, Braun KR, Van Biber BP, Laflamme MA, Murry CE and Wight TN. Differentiation of cardiomyocytes from human embryonic stem cells is accompanied by changes in the extracellular matrix production of versican and hyaluronan. J Cell Biochem. 2010;111:585–96.

57. Hang CT, Yang J, Han P, Cheng HL, Shang C, Ashley E, Zhou B and Chang CP. Chromatin regulation by Brg1 underlies heart muscle development and disease. Nature. 2010;466:62–7.

58. Hota SK, Johnson JR, Verschueren E, Thomas R, Blotnick AM, Zhu Y, Sun X, Pennacchio LA, Krogan NJ and Bruneau BG. Dynamic BAF chromatin remodeling complex subunit inclusion promotes temporally distinct gene expression programs in cardiogenesis. Development. 2019;146.

59. Singh AP, Foley JF, Rubino M, Boyle MC, Tandon A, Shah R and Archer TK. Brg1 Enables Rapid Growth of the Early Embryo by Suppressing Genes That Regulate Apoptosis and Cell Growth Arrest. Mol Cell Biol. 2016;36:1990–2010.

60. Shimono Y, Murakami H, Kawai K, Wade PA, Shimokata K and Takahashi M. Mi-2 beta associates with BRG1 and RET finger protein at the distinct regions with transcriptional activating and repressing abilities. J Biol Chem. 2003;278:51638–45.

61. Sun X, Yu W, Li L and Sun Y. ADNP Controls Gene Expression Through Local Chromatin Architecture by Association With BRG1 and CHD4. Front Cell Dev Biol. 2020;8:553.

62. Morris SA, Baek S, Sung MH, John S, Wiench M, Johnson TA, Schiltz RL and Hager GL. Overlapping chromatin-remodeling systems collaborate genome wide at dynamic chromatin transitions. Nat Struct Mol Biol. 2014;21:73–81.

63. Dyer LA and Patterson C. A novel ex vivo culture method for the embryonic mouse heart. J Vis Exp. 2013:e50359.

64. Cao J and Poss KD. Explant culture of adult zebrafish hearts for epicardial regeneration studies. Nat Protoc. 2016;11:872–81.

65. Rhee S, Chung JI, King DA, D’Amato G, Paik DT, Duan A, Chang A, Nagelberg D, Sharma B, Jeong Y, Diehn M, Wu JC, Morrison AJ and Red-Horse K. Endothelial deletion of Ino80 disrupts coronary angiogenesis and causes congenital heart disease. Nat Commun. 2018;9:368.

66. Powers N and Huang GN. Visualization of regenerating and repairing hearts. Clin Sci (Lond). 2022;136:787–798.

67. Rhee S, Paik DT, Yang JY, Nagelberg D, Williams I, Tian L, Roth R, Chandy M, Ban J, Belbachir N, Kim S, Zhang H, Phansalkar R, Wong KM, King DA, Valdez C, Winn VD, Morrison AJ, Wu JC and Red-Horse K. Endocardial/endothelial angiocrines regulate cardiomyocyte development and maturation and induce features of ventricular non-compaction. Eur Heart J. 2021;42:4264–4276.

68. Ham SA, Yoo T, Lee WJ, Hwang JS, Hur J, Paek KS, Lim DS, Han SG, Lee CH and Seo HG. ADAMTS1-mediated targeting of TSP-1 by PPARdelta suppresses migration and invasion of breast cancer cells. Oncotarget. 2017;8:94091–94103.

69. Pinard A, Guey S, Guo D, Cecchi AC, Kharas N, Wallace S, Regalado ES, Hostetler EM, Sharrief AZ, Bergametti F, Kossorotoff M, Herve D, Kraemer M, Bamshad MJ, Nickerson DA, Smith ER, Tournier-Lasserve E and Milewicz DM. The pleiotropy associated with de novo variants in CHD4, CNOT3, and SETD5 extends to moyamoya angiopathy. Genet Med. 2020;22:427-431.

70. Sifrim A, Hitz MP, Wilsdon A, Breckpot J, Turki SH, Thienpont B, McRae J, Fitzgerald TW, Singh T, Swaminathan GJ, Prigmore E, Rajan D, Abdul-Khaliq H, Banka S, Bauer UM, Bentham J, Berger F, Bhattacharya S, Bu’Lock F, Canham N, Colgiu IG, Cosgrove C, Cox H, Daehnert I, Daly A, Danesh J, Fryer A, Gewillig M, Hobson E, Hoff K, Homfray T, Study I, Kahlert AK, Ketley A, Kramer HH, Lachlan K, Lampe AK, Louw JJ, Manickara AK, Manase D, McCarthy KP, Metcalfe K, Moore C, Newbury-Ecob R, Omer SO, Ouwehand WH, Park SM, Parker MJ, Pickardt T, Pollard MO, Robert L, Roberts DJ, Sambrook J, Setchfield K, Stiller B, Thornborough C, Toka O, Watkins H, Williams D, Wright M, Mital S, Daubeney PE, Keavney B, Goodship J, Consortium UK, Abu-Sulaiman RM, Klaassen S, Wright CF, Firth HV, Barrett JC, Devriendt K, FitzPatrick DR, Brook JD, Deciphering Developmental Disorders S and Hurles ME. Distinct genetic architectures for syndromic and nonsyndromic congenital heart defects identified by exome sequencing. Nat Genet. 2016;48:1060–5.

71. Tian X, Li Y, He L, Zhang H, Huang X, Liu Q, Pu W, Zhang L, Li Y, Zhao H, Wang Z, Zhu J, Nie Y, Hu S, Sedmera D, Zhong TP, Yu Y, Zhang L, Yan Y, Qiao Z, Wang QD, Wu SM, Pu WT, Anderson RH and Zhou B. Identification of a hybrid myocardial zone in the mammalian heart after birth. Nat Commun. 2017;8:87.

72. Wu T, Liang Z, Zhang Z, Liu C, Zhang L, Gu Y, Peterson KL, Evans SM, Fu XD and Chen J. PRDM16 Is a Compact Myocardium-Enriched Transcription Factor Required to Maintain Compact Myocardial Cardiomyocyte Identity in Left Ventricle. Circulation. 2022;145:586–602.

73. Choquet C, Nguyen THM, Sicard P, Buttigieg E, Tran TT, Kober F, Varlet I, Sturny R, Costa MW, Harvey RP, Nguyen C, Rihet P, Richard S, Bernard M, Kelly RG, Lalevee N and Miquerol L. Deletion of Nkx2-5 in trabecular myocardium reveals the developmental origins of pathological heterogeneity associated with ventricular non-compaction cardiomyopathy. PLoS Genet. 2018;14:e1007502.

74. Chen H, Zhang W, Sun X, Yoshimoto M, Chen Z, Zhu W, Liu J, Shen Y, Yong W, Li D, Zhang J, Lin Y, Li B, VanDusen NJ, Snider P, Schwartz RJ, Conway SJ, Field LJ, Yoder MC, Firulli AB, Carlesso N, Towbin JA and Shou W. Fkbp1a controls ventricular myocardium trabeculation and compaction by regulating endocardial Notch1 activity. Development. 2013;140:1946–57.

75. Priya R, Allanki S, Gentile A, Mansingh S, Uribe V, Maischein HM and Stainier DYR. Tension heterogeneity directs form and fate to pattern the myocardial wall. Nature. 2020;588:130–134.

76. Iruela-Arispe ML, Carpizo D and Luque A. ADAMTS1: a matrix metalloprotease with angioinhibitory properties. Ann N Y Acad Sci. 2003;995:183–90.

77. Venkatesh DA, Park KS, Harrington A, Miceli-Libby L, Yoon JK and Liaw L. Cardiovascular and hematopoietic defects associated with Notch1 activation in embryonic Tie2-expressing populations. Circ Res. 2008;103:423–31.

78. Yang J, Bucker S, Jungblut B, Bottger T, Cinnamon Y, Tchorz J, Muller M, Bettler B, Harvey R, Sun QY, Schneider A and Braun T. Inhibition of Notch2 by Numb/Numblike controls myocardial compaction in the heart. Cardiovasc Res. 2012;96:276–85.

79. Watanabe Y, Kokubo H, Miyagawa-Tomita S, Endo M, Igarashi K, Aisaki K, Kanno J and Saga Y. Activation of Notch1 signaling in cardiogenic mesoderm induces abnormal heart morphogenesis in mouse. Development. 2006;133:1625–34.

80. Mysliwiec MR, Bresnick EH and Lee Y. Endothelial Jarid2/Jumonji is required for normal cardiac development and proper Notch1 expression. J Biol Chem. 2011;286:17193–204.

81. Jan-Renier A.J. Moonen JC, Minyi Shi, Tsutomu Shinohara, Dan Li, Maxwell R. Mumbach, Fan Zhang, Joseph Nasser, Daniel H. Mai, Shalina Taylor, Lingli Wang, Ross J. Metzger, Howard Y. Chang, Jesse M. Engreitz, Michael P. Snyder, Marlene Rabinovitch. KLF4 Recruits SWI/SNF to Increase Chromatin Accessibility and Reprogram the Endothelial Enhancer Landscape under Laminar Shear Stress. BioRxiv. 2020.

## Supplemental References

1. Robbe ZL, Shi W, Wasson LK, Scialdone AP, Wilczewski CM, Sheng X, Hepperla AJ, Akerberg BN, Pu WT, Cristea IM, Davis IJ and Conlon FL. CHD4 is recruited by GATA4 and NKX2-5 to repress noncardiac gene programs in the developing heart. Genes Dev. 2022;36:468–482.

2. Wilczewski CM, Hepperla AJ, Shimbo T, Wasson L, Robbe ZL, Davis IJ, Wade PA and Conlon FL. CHD4 and the NuRD complex directly control cardiac sarcomere formation. Proc Natl Acad Sci U S A. 2018;115:6727–6732.

3. Stankunas K, Hang CT, Tsun ZY, Chen H, Lee NV, Wu JI, Shang C, Bayle JH, Shou W, Iruela-Arispe ML and Chang CP. Endocardial Brg1 represses ADAMTS1 to maintain the microenvironment for myocardial morphogenesis. Dev Cell. 2008;14:298–311.

4. Dorr KM, Amin NM, Kuchenbrod LM, Labiner H, Charpentier MS, Pevny LH, Wessels A and Conlon FL. Casz1 is required for cardiomyocyte G1-to-S phase progression during mammalian cardiac development. Development. 2015;142:2037–47.

5. Cao J and Poss KD. Explant culture of adult zebrafish hearts for epicardial regeneration studies. Nat Protoc. 2016;11:872–81.

6. Dyer LA and Patterson C. A novel ex vivo culture method for the embryonic mouse heart. J Vis Exp. 2013:e50359.

7. Ehler E, Moore-Morris T and Lange S. Isolation and culture of neonatal mouse cardiomyocytes. J Vis Exp. 2013.

8. Mei L, Kedziora KM, Song EA, Purvis JE and Cook JG. The consequences of differential origin licensing dynamics in distinct chromatin environments. Nucleic Acids Res. 2022.

9. Waldron L, Steimle JD, Greco TM, Gomez NC, Dorr KM, Kweon J, Temple B, Yang XH, Wilczewski CM, Davis IJ, Cristea IM, Moskowitz IP and Conlon FL. The Cardiac TBX5 Interactome Reveals a Chromatin Remodeling Network Essential for Cardiac Septation. Dev Cell. 2016;36:262–75.

10. Shi W, Sheng X, Dorr KM, Hutton JE, Emerson JI, Davies HA, Andrade TD, Wasson LK, Greco TM, Hashimoto Y, Federspiel JD, Robbe ZL, Chen X, Arnold AP, Cristea IM and Conlon FL. Cardiac proteomics reveals sex chromosome-dependent differences between males and females that arise prior to gonad formation. Dev Cell. 2021;56:3019–3034 e7.

11. Kim GH. Murine fetal echocardiography. J Vis Exp. 2013.

12. Kim W, Seidah NG and Prat A. In utero measurement of heart rate in mouse by noninvasive M-mode echocardiography. J Vis Exp. 2013:e50994.

13. Touma M. Fetal Mouse Cardiovascular Imaging Using a High-frequency Ultrasound (30/45MHZ) System. J Vis Exp. 2018.

14. Mootha VK, Lindgren CM, Eriksson KF, Subramanian A, Sihag S, Lehar J, Puigserver P, Carlsson E, Ridderstrale M, Laurila E, Houstis N, Daly MJ, Patterson N, Mesirov JP, Golub TR, Tamayo P, Spiegelman B, Lander ES, Hirschhorn JN, Altshuler D and Groop LC. PGC-1alpha-responsive genes involved in oxidative phosphorylation are coordinately downregulated in human diabetes. Nat Genet. 2003;34:267–73.

15. Subramanian A, Tamayo P, Mootha VK, Mukherjee S, Ebert BL, Gillette MA, Paulovich A, Pomeroy SL, Golub TR, Lander ES and Mesirov JP. Gene set enrichment analysis: a knowledge-based approach for interpreting genome-wide expression profiles. Proc Natl Acad Sci U S A. 2005;102:15545–50.

